# PAM-less Exonuclease-assisted Cas12a for visual detection of Vibrio Species

**DOI:** 10.1101/2022.10.21.513145

**Authors:** Derek Han Zhang, Siddharth Raykar, Kenneth Tsz Chun Ng

## Abstract

Foodborne pathogens, including *Vibrio spp*. and norovirus, cause substantial economic and healthcare burdens worldwide. Rapid and sensitive point-of-care testing on-farm or restaurants for batch inspection of pathogenic contamination in raw food products is essential. Here, we present an easy-to-design, cost-effective PAM-less Exonuclease-assisted Cas12A Nucleic-acid Detection (PECAN) assay paired with nucleic acid amplification systems for rapid and sensitive visual detection of 2 pathogenic Vibrio species: *Vibrio parahaemolyticus* (*TDH*) and *Vibrio Cholerae* (*ctxA*) without protospacer adjacent motif (PAM) site limitation. With T7 exonuclease, PAM-less detection could be achieved with a low concentration of cas12a, costing $0.8 USD per reaction. The system could also be adapted for PAM-less cas12a nucleic acid detection in-field or in-lab for sensitive DNA or RNA detection. We also constructed a low-cost reusable 3D printed heater chassis and reusable sodium acetate heat packs for field use without generating solid waste.

## Introduction

Global warming has increased the proliferation of pathogenic bacterial species in water [1,2]. It was estimated that they bring an economic impact of 7 billion USD in the United States alone every year. Children under five, the immunocompromised and the elderly contributing to most cases of foodborne illness-related deaths. Thus, there is an urgent need for new tools to detect foodborne pathogens [3].

*Vibrio parahaemolyticus* is one of the most common causes of foodborne illness worldwide [4,5], causing acute gastroenteritis and complications. *Vibrio Cholerae* is an infectious agent which causes Cholera [6]. Consumption of raw or undercooked contaminated seafood may lead to infections, and shellfish contribute to a large proportion of Vibrio infections due to their filter feeder ability which leads to the accumulation of as much as a 100-fold bacterial concentration within the organism [6]. In our study, to identify virulent Vibrio strains, thermostable direct hemolysin (*TDH*) for *Vibrio parahaemolyticus* [7] and Cholera Toxin Genes (*ctxA*) for *Vibrio Cholerae* [8] were chosen as the detection markers. Current commercialised Vibrio detection includes qPCR kits, or relies on biochemical properties such as fermentation and selective agar for visualising colony morphology and colourimetric display.

Clustered regularly interspaced short palindromic repeats (CRISPR) enzymes, such as those within the cas12 and cas13 family, were used to detect nucleic acid with high specificity and sensitivity [7–9]. CRISPR enzymes are activated upon nucleic acid recognition by crRNA hybridisation and exhibit non-specific single-stranded DNA (ssDNA) cleavage activity. By incorporating an ssDNA reporter activated through cleavage of the ssDNA linker between the fluorophore and quencher (ssDNA-FQ), which leads to a visual fluorescence signal. The use of CRISPR detection has often been paired with an exponential nucleic acid amplification methods, including polymerase chain reaction (PCR) or isothermal amplification, including recombinase polymerase amplification (RPA) and loop-mediated isothermal amplification (LAMP). While using nucleic acid amplifications as the sole method of detection might lead to false positives [10,11], the incorporation of CRISPR enzymes allows increased specificity of detection. These methods have been proven successful in achieving sensitive detection of viruses [12–14], bacterial species [15,16] or coupled with aptamers for molecular detection [17,18].

Cas12a ssDNA-FQ trans-cleavage activity could be activated by cas12a-crRNA complex hybridizing to template ssDNA, or be activated by double-stranded DNA (dsDNA) with a protospacer adjacent motif (PAM) site, but not ssRNA templates [8]. Traditionally, cas12a detection methods were designed to recognise the PAM sequence of TTTV with a double-stranded template. Upon binding to the PAM sequence, the Cas12a protein unwound the dsDNA at the site of interest and allowed for crRNA hybridisation and cas12a activation [19]. However, the PAM site might not always be present within the target of interest or conserved region of the detection site. Hence, researchers have investigated other strategies to deal with the PAM site limitation to allow for increased flexibility in detection design, via enzyme engineering [20,21] and exploring orthologs [22–24] to increase the suite of PAM sites available for cas12a recognition and activation.

Cas12a detection without PAM activation has also been published previously. Some previous strategies include LAMP amplification pairing with FnCas12a (Cas-PfLAMP) [25], PAM-less conditional DNA substrate [26], and All-In-One Dual CRISPR-Cas12a (AIOD-CRISPR). However, not all these are appropriate for *in vitro* nucleic detection for all use cases. For example, LAMP isothermal amplification requires a high activation temperature between 60 and 65°C [27], which might not apply to field detection. PAM-less conditional DNA substrate [26] requires mismatches to generate a pre-wound site for bubble generation, which increases the complexity in primer design and amplification steps. While the AIOD-CRISPR system were used for ultrasensitive and rapid point-of-care detection of coronavirus SARS-CoV-2. They utilised the ssDNA generated from the RPA strand-displacing recombinase, which hybridises to cas12a crRNA, activating the cas12a trans-cleavage activity for ssDNA-FQ cleavage to generate a visual output [13]. Their research paves the way for our system, demonstrating the use of ssDNA as the cas12a activator for rapid, specific and sensitive output. However, the AIOD-CRISPR system uses a very high concentration of cas12a (1280nM), and with another group optimising the protocol, the usage of cas12a is still high (1000nM) [28]. In comparison, RPA-cas12a detection utilizing the PAM site generally utilize cas12a concentration of 30nM to 200nM [29–33]. The underlying reason for the usage of such high cas12a concentration in AIOD-CRISPR is due to buffer incompatibility, as cas12a activity is drastically reduced in RPA buffer. The buffer incompatibility issue also led other groups to develop the dynamic aqueous multiphase reaction (DAMR), which achieved more than a hundred times fluorescence compared to utilizing cas12a within RPA buffer [34].

Moreover, a higher concentration of cas12a was reported to lead to decreased specificity of ssDNA template recognition. It was shown that higher concentrations of cas12a are activated by shorter ssDNA sequences, compared to no activation by shorter ssDNA with a lower concentration of cas12a, even with longer incubation time [35]. Hence there might be higher false positive rates of detection in the case when there is a partial crRNA complementary sequence in the RPA amplicon. Additionally, the use of high concentrations of cas12a increases the cost of detection multiple folds, thereby decreasing the willingness of the food industry and individual restaurant owners to carry out routine batch-based detection of their food products. Therefore, it would be desirable to construct a cas12a ssDNA-based detection assay with lower cas12a concentration and optimised buffering.

Here, we present the PAM-less Exonuclease-assisted Cas12A Nucleic-acid Detection (PECAN) system. The system utilises T7 exonuclease (T7exo), a 5’ to 3’ DNA exonuclease that recognises dsDNA at blunt or nicked sites [36]. T7exo was utilized in previous study to produce ssDNA from RPA reaction for lateral flow detection of SARS-CoV-2 [37]. T7exo does not cleave ssDNA or ssRNA and was proven for efficient ssDNA production *in vitro* using phosphorothioate-protected DNA primers[38], in which phosphorothioate bond was shown to inhibit T7exo digestion initiation at one strand of the dsDNA. Our study presents an optimised RPA/Cas12a/T7exo system for sensitive detection of Vibrio species, with a cost per reaction of $0.8 USD. We also demonstrated the system’s versatility, showing that PECAN detection applies not only to the RPA one-pot assay but applicable to other amplification methods such as PCR for dual validation.

## Results

### The PECAN system activates Cas12a trans-cleavage without the PAM site

Conventionally, to activate the cas12a trans-cleavage with dsDNA substrate, a PAM site is required for cas12a to unwind the dsDNA to create a bubble (triple-strand DNA-RNA R-loop) for complementary site binding [38]. A canonical PAM site is TTTV for Lbcas12a, while other non-canonical PAM could also lead to cas12a activation, albeit with lower activity, for example, CTTV, TCTV and TTCV [39]. The crRNA-template hybridisation causes a conformation change to the REC lobe, leading to activation of the finger domain and RuvC-NuC pocket catalytic activity. While a PAM site is required for the bubble formation, it does not contribute to the catalytic activation. Both dsDNA and ssDNA template requires a crRNA complementary region of at least 14nt to activate the trans-cleavage activity, showing that the crRNA +15 position are critical to cas12a activation and that it is PAM independent [38]. Therefore, ssDNA could also be used as a template for cas12a trans-cleavage activation. To test the PAM-less activation of cas12a trans-cleavage activity with PECAN, we designed two crRNA probes without PAM sites for the visual detection of *Vibrio parahaemolyticus TDH* and *Vibrio Cholerae ctxA* genes. Instead of searching and screening for crRNAs with PAM sites (TTTV), we directly designed the crRNA sequence with conserved regions, with a template site preceding the 5’ end crRNA being CATG (*TDH*) and GTCT (*ctxA*).

To utilise cas12a-mediated activation without PAM, dsDNA is first digested to form ssDNA templates containing the complementary sequence to the crRNA. While there are other ssDNA exonucleases that could generate ssDNA from dsDNA templates, i.e., 3’ to 5’ exonucleases like exonuclease I, these cannot be utilised since it is hard to build 3’ DNA protection due to the RPA nature of 5’ to 3’ extension. 5’ to 3’ exonucleases which do not exhibit untagged ssDNA digestion activity include Lambda exonuclease [39], Exonuclease VIII [40] and T7exo. T7exo was well characterised in its ability to generate ssDNA from dsDNA efficiently. In comparison, Lambda exonuclease requires HPLC-grade phosphorylated primers and often results in incomplete digestion [38]. Therefore, T7exo was chosen to be tested for constructing the PECAN system.

T7exo 5’ to 3’ end digestion was blocked by replacing the DNA phosphodiester bond of 5-6nt at the 5’ with phosphorothioate bonds, which was included the 5’ end of the primers [41]. T7exo would digest only the complementary strand, exposing the crRNA binding site on the template strand. Therefore, PECAN is a one-pot system, with T7exo providing ssDNA templates for cas12a activation.

In Fig. 1b, fluorescence was not observed in controls (no template, RPA reaction core, or cas12a), showing that the reagents did not cause cas12a-mediated trans-cleavage, but showing high fluorescence signal when the DNA amplicons from RPA were present. Moreover, with a plasmid with a TTTA PAM site added before the *TDH*1 crRNA, the activation of cas12a could be observed without T7exo, but not with the template without the PAM site. However with the addition of T7exo, fluorescence could be observed from template with or without PAM site (Supplementary Fig. 13). There was weak fluorescence in the ‘No T7exo’ control, in which small amount of ssDNA naturally occur in the RPA reaction. The AIOD-CRISPR system [13] utilises RPA strand displacements to provide ssDNA for PAM-less activation of cas12a. The strand displacement activity of RPA in PECAN without T7exo might have caused the activation of cas12a, albeit with much lower fluorescence. Lastly, a high fluorescence was observed with every PECAN component utilised.

**Fig. 1.**
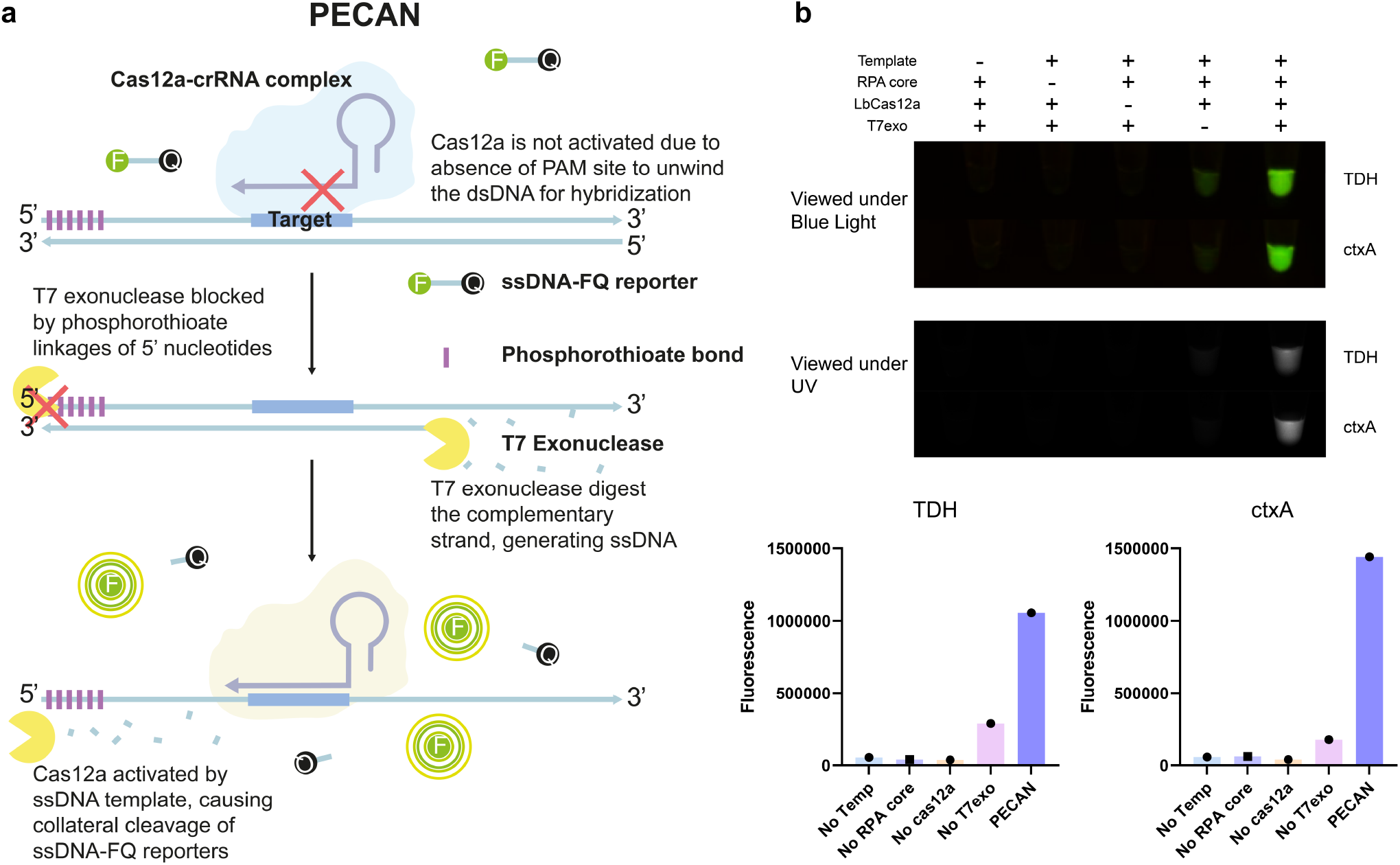
Design and working principle of PECAN assay. **a** Schematic of the PECAN system. The cas12a-crRNA complex is complementary to the target, the cross denotes the block of hybridisation without a PAM site mediated bubble. **b** Evaluation of 5 different PECAN reactions at 20-min endpoints under blue or UV light. The ssDNA-FQ reporter sequence is /56-FAM/TTATT/3BHQ_1/ with 5’ Fluorescein and 3’ Black Hole Quencher 1 modification, which produces fluorescence upon cleavage of ssDNA for evaluation of cas12a trans-cleavage activity. PECAN compose of 0.1unit μL^-1^ of T7exo, 8μM ssDNA-FQ, 0.2μM EnGen Lba Cas12a, with 0.4μM crRNA. crRNA *TDH*: *TDH*1_3’7DNA, for *ctxA*: *ctxA*_R2 (Supplementary Table 1). The reaction was carried out in “Mix” buffer (0.5x NEB 2.1 + 0.5x NEB 4) with a final DTT concentration of 1.5mM DTT. A plasmid containing *TDH* and *ctxA* gene sequence (3 × 10^3^ copies) was used in a 10μL RPA reaction, in which 0.5μL of the RPA reaction (20 minutes) was added to the PECAN mix.

### crRNA and primer design based on bioinformatics analysis

False negative readouts could be prevented with crRNA target locating at highly conserved regions of the genome where mutations are unlikely. After identifying a pathogenic marker, for example, *TDH* for *Vibrio parahaemolyticus*, a multiple sequence alignment was ran to identify the regions with consensus (Supplementary Table 6). crRNA could then be designed on a conserved region. Although RPA primers are more tolerant to single base pair mismatches [42], care should still be taken to locate a conserved region to minimise the use of degenerate primers.

6-nt phosphorothioate site modification was added to primers at the 5’ end. The crRNA *TDH*1 used for Fig.2 is a forward-direction crRNA, and the target site is amplified with the reverse primer, therefore PECAN would be efficient with a reverse primer phosphorothioate protection. For amplicons without any phosphorothioate protection, T7exo could still initiate and digest both strands of the dsDNA, leaving ssDNA on either side. While this would also trigger the activation of cas12a, the digested ssDNA may re-hybridise and trigger another round of T7exo digestion. However, if the template amount is in excess compared to available cas12a, non-protected primers might be used as an alternative to phosphorothioate-protected primers to trigger ssDNA-mediated cas12a activation. For amplicons with phosphorothioate protected forward primers, the reverse strand of the amplicon would be digested, leading to the loss of the crRNA hybridisation site. While for amplicons with protected forward and reverse primer, both sides would be resistant to T7exo digestion, and hence the target strand is not exposed to trigger crRNA hybridisation. Consistent to the design, only the amplicons with protected reverse primer have a strong signal due to the exposed reverse target strand to the forward direction crRNA. It demonstrated that PECAN was efficient with the correct combination of crRNA and phosphorothioate primer.

**Fig. 2.**
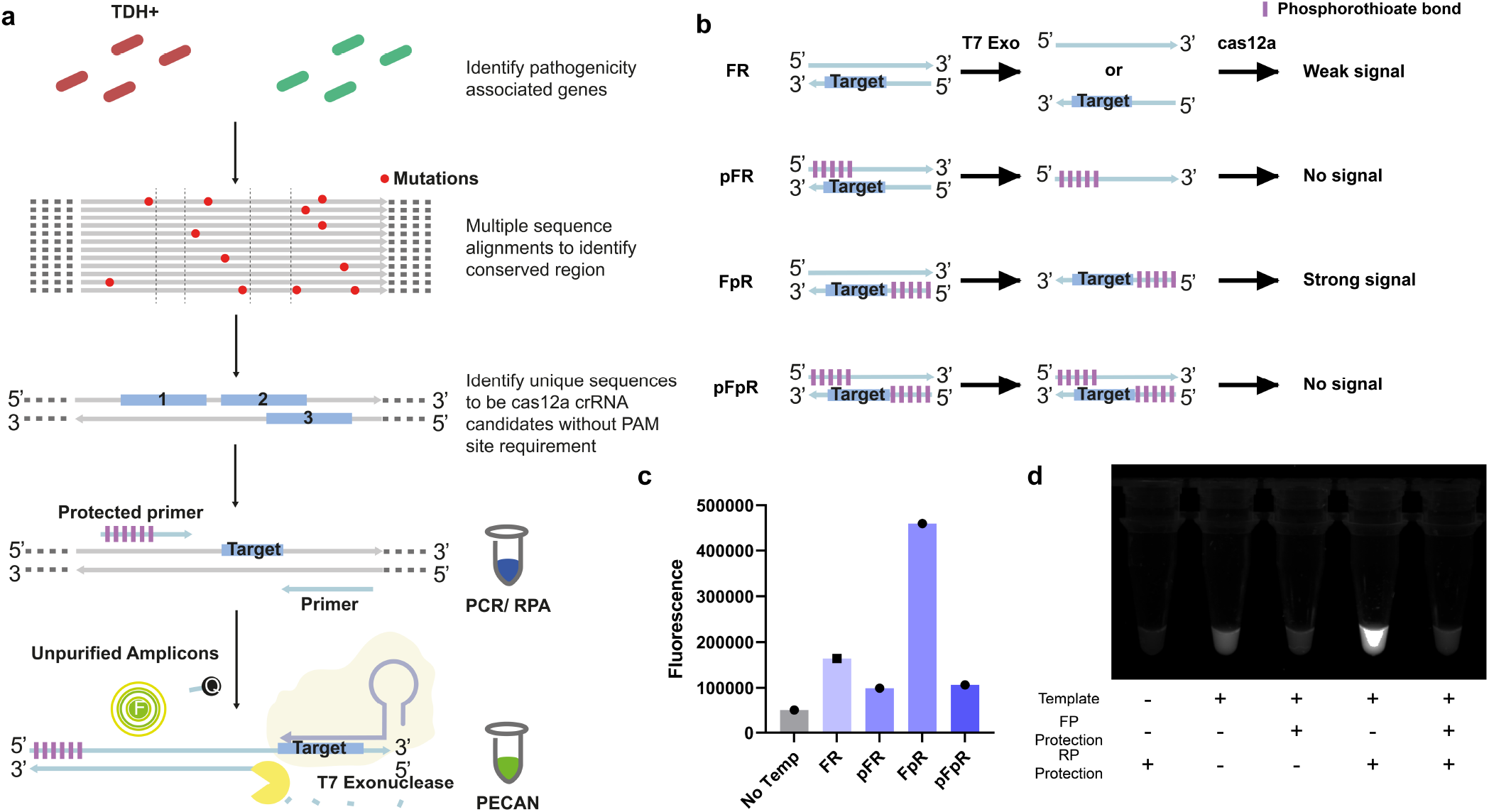
Example for designing crRNA and primers for PECAN detection. **a** Workflow of identifying appropriate regions for crRNA and primer design. **b** The inclusion of phosphorothioate protection at different primers produces different outcomes of PECAN. **c** Fluorescence measurement of 20-min PECAN combinations of protected RPA primers for *TDH* (Supplementary Table 3) and crRNA *TDH*1 (Supplementary Table 1). **d** UV view of 20-min PECAN with different combinations of protected RPA primers for *TDH*, with crRNA *TDH*1.

### Optimisation of T7exo and cas12a for PECAN

NEB recommended NEB r2.1 buffer for EnGen Lba cas12a and NEB 4 buffer for T7exo. For initial screening for buffer optimal for T7exo and cas12a activity for the one-pot system, various buffers, including NEB r2.1, Mix (0.5x NEB r2.1 + 0.5x NEB 4), and NEB 4, were tested with various DTT and Mg2+ concentration. For T7exo, it was found that consistent with NEB reports, T7exo lost most of the activity in NEB r2.1 buffer (Supplementary Fig. 4). While comparing T7exo in NEB 4 or Mix, T7exo activity is enhanced in Mix although having a lower DTT concentration in Mix. With the addition of DTT to a final of 1mM, T7exo activity is further enhanced. However, with the Addition of MgCl_2_ to Mg^2+^ concentration of 20mM, T7exo activity is hindered (Fig. 3a). It indicates that the addition of DTT provides a good reducing environment for T7exo to be active. While further increasing the DTT concentration from 1mM to 40mM in Mix buffer did not make further enhancement of T7exo activity (Fig. 3c). While comparing cas12a in various buffers, it was found that cas12a maintained the cleavage activity in NEB 4 and Mix compared to NEB r2.1 (Fig. 3b). Addition to 1mM of DTT did not improve cas12a activity significantly, however, with 10mM of DTT, cas12a trans-cleavage activity increased (Fig. 3b, d, Supplementary Fig 2-3). It is worth noting that the addition of DTT increased the background of no template control for crRNA *TDH*1 (Supplementary Fig. 7) but not in *TDH*1_3’7DNA (Supplementary Fig. 2), possibly due to the remaining undigested ssDNA template of crRNA *TDH*1 from *in vitro* transcription reaction, or some background cleavage mechanism blocked by crRNA 3’7DNA modification. It was also shown that increasing magnesium concentration to 20mM with added DTT could enhance cas12a trans-cleavage activity. However, since added magnesium inhibited T7exo activity in both Mix and Mix+DTT (Fig. 3a), the optimal buffer for one-pot cas12a-T7exo would be Mix+DTT.

**Fig. 3.**
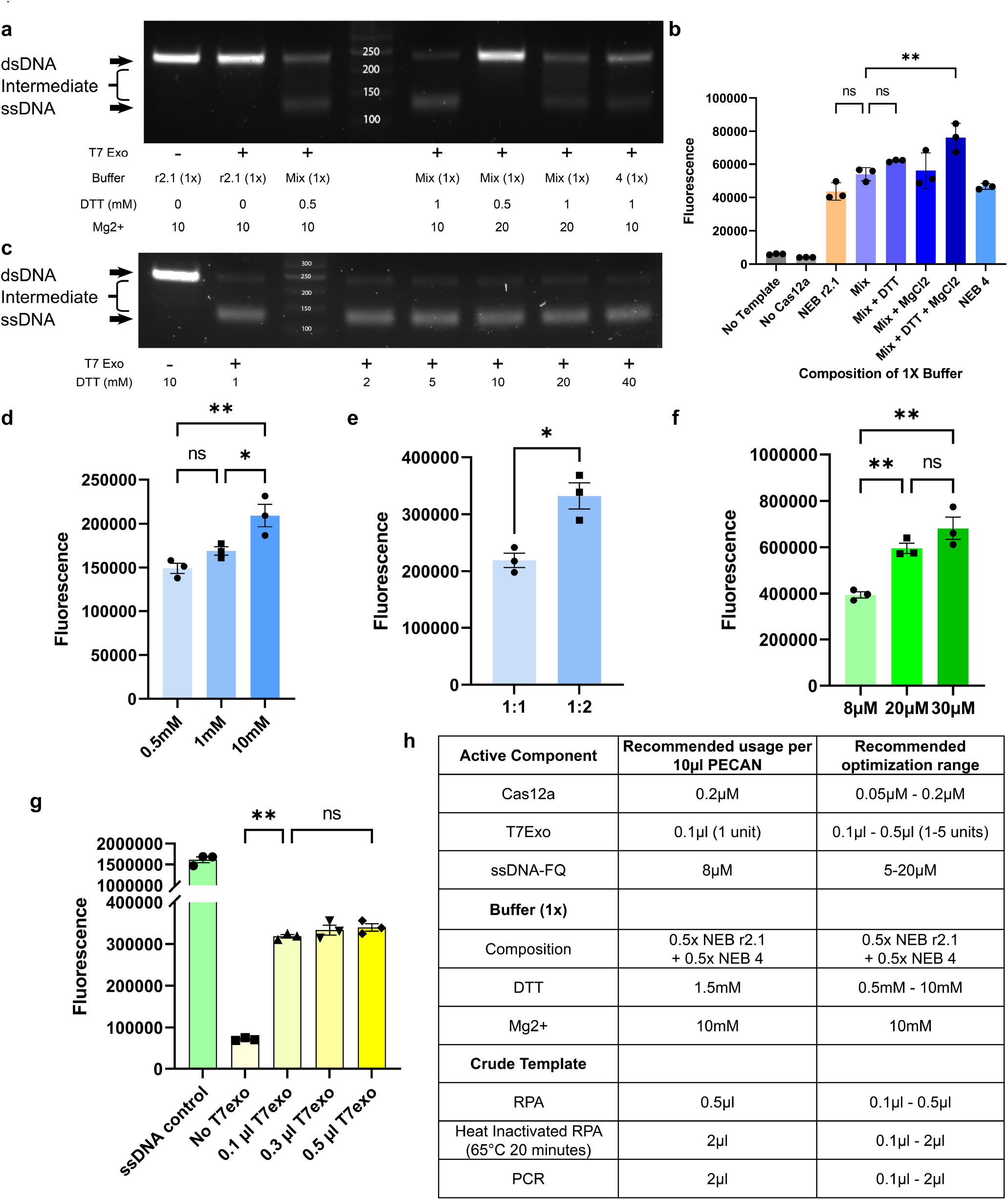
Optimisation of T7exo and cas12a one-pot system. **a** T7exo buffer optimisation with various buffering conditions, DTT and Mg^2+^ concentration (Supplementary Fig. 4). **b** cas12a buffer optimisation with various buffering conditions, DTT and Mg^2+^ concentration. **c** T7exo DTT optimisation above 1mM. **d** cas12a DTT optimisation with 10x molar ssDNA as a template. **e** transcleavage activity observed for different cas12a: crRNA ratios, after 10-min room temperature cas12a-crRNA coupling. **f** ssDNA-FQ concentration optimization for cas12a trans-cleavage. **g** PECAN T7exo amount optimisation. **h** Recommended reagent concentration, buffer composition and template ratio for PECAN. 0.1μM cas12a, 0.2μM crRNA *TDH*1 and 1μM of ssDNA template were used for **b, d, e, f**. With 1-unit T7exo was used for **a, c** with 100ng of *TDH* amplicon in 10μL reaction, 20 min incubation. Error bars represent the means ± S.E.M. from replicates. The original one-way ANOVA was used to analyse the statistical significance of **b, d, f, g**. The unpaired two-tailed t-test was used to analyse the statistical significance of **e**.

**Fig. 4.**
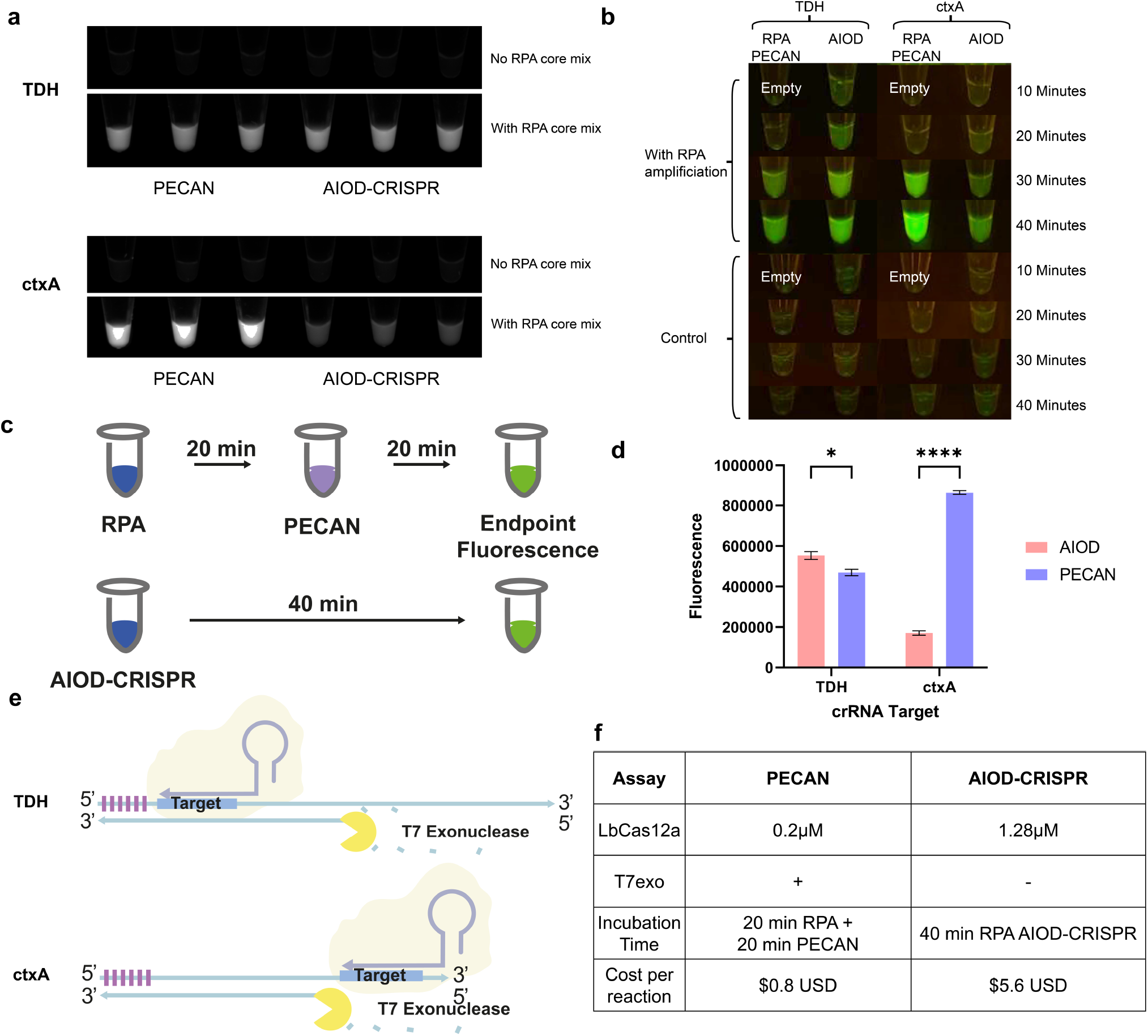
Comparison between PECAN and AIOD-CRISPR. **a** UV view of PECAN and AIOD-CRISPR at the 40-min endpoint to detect the presence of *TDH* and *ctxA* DNA with or without RPA core reaction. **b** Blue light view of PECAN and AIOD-CRISPR at 10-40 min with or without RPA core reaction. **c** Schematics of the PECAN and AIOD-CRISPR, for PECAN, the RPA was first incubated 20-min, and 0.5μL of RPA reaction was transferred to the PECAN mix with 0.2μM of cas12a and 1 unit of T7exo, followed by 20-min incubation for a 40-min endpoint measurement. The same RPA mix was used in AIOD-CRISPR, and a final of 1.28μM of cas12a was added to the reaction mix. The cas12a-crRNA incubation was done for 10-minutes at room temperature before the start, and the same mix was used to assemble AIOD-CRISPR and PECAN. The PECAN mix with cas12a was was put on ice during the first 20-min of the experiment. The plasmid template used was 3×10^3^ per 10μL of RPA reaction. **d** Fluorescence measurement of *TDH* and *ctxA* detection at the 40-minute endpoint. **e** crRNA position of *TDH*1 and *ctxA*_R2 relative to the primers within the amplicon. **f** Table for comparison of PECAN and AIOD-CRISPR in cas12a, T7exo usage and cost per reaction.

For a rapid 10-min room temperature incubation of cas12a with crRNA, a 1:2 ratio for cas12a:crRNA was found to yield higher trans-cleavage activity compared to a 1:1 ratio (Fig. 3e, Supplementary Fig. 8). For fluorophore optimisation, it was found that the fluorescence signal for measuring cas12a trans-cleavage activity peaked at 20-30μM of fluorophore (Supplementary Fig. 5), while there was a significant increase of fluorescence of using 20μM of ssDNA-FQ compared to 8μM (Fig. 3f). However, the improvement is less than 2-fold, and will lead to increased cost of detection. Using a higher fluorophore concentration also resulted in higher background fluorescence (Supplementary Fig. 6). For crRNA, a dual crRNA method was also tested. With crRNA *TDH*1 and *TDH*2, when combined, yielded a higher fluorescence with the same final crRNA concentration, possibly due to the doubling of available sites for crRNA binding per strand of ssDNA template. However, the improvement in signal for *TDH*1 and TDH2 is less than 2-fold compared to using *TDH*1 or *TDH*2 separately (Supplementary Fig. 10). The extent of improvement with dual crRNA might also be relative to the total template available compared to the total cas12a amount in the solution.

While optimising T7 usage, it was observed that with the direct mixing of RPA with the PECAN system, an equilibrium position existed, indicating that RPA was still active within the PECAN mix. This RPA activity converted ssDNA generated via T7exo digestion back to dsDNA (Supplementary Figure 12). The equilibrium caused the decrease of fluorescence due to the competition of ssDNA between RPA enzymes and cas12a. However, decreasing the RPA volume transferred to PECAN can shift the equilibrium position from dsDNA to ssDNA in RPA-PECAN. The equilibrium of ssDNA and dsDNA was not observed with PCR samples (Supplementary Fig. 12). To address the question of whether the equilibrium position is dominated by RPA or T7exo, with the same amount of RPA solution added to PECAN, decreasing the usage of T7exo did not lead to the shift of ssDNA position back to dsDNA position, indicating that T7exo is in excess. The equilibrium position is maintained by the ratio of RPA to PECAN. It was verified in agarose gel (Supplementary Fig. 9), and in PECAN with decreased amount of T7exo added (Fig. 3g), with no significant difference in fluorescence observed. Furthermore, the shifting of equilibrium from ssDNA to dsDNA by RPA enzymes was broken with heat denaturation of RPA solution before being added to PECAN. Interestingly, only a 20-min 65°C heat inactivation would result in the complete conversion of dsDNA to ssDNA by T7exo but not 85°C or 100°C, and the result was also consistent in PECAN with a high fluorescence observed when RPA has undergone a 20-min 65°C heat inactivation (Supplementary Figure 11).

### Comparison between PECAN with previous studies

When comparing PECAN and AIOD-CRISPR, it was observed that within the same 40-min timeframe, the fluorescence produced for detecting *TDH* with AIOD-CRISPR assay was slightly higher than PECAN. While for detecting *ctxA*, the fluorescence of PECAN is >4-fold higher than the AIOD-CRISPR assay. The PECAN is a robust system that can achieve similar or enhanced signal development with the same starting template compared to AIOD-CRISPR, with >6 fold less cas12a usage without an increase in background fluorescence (Supplementary Fig. 14).

While considering the discrepancy between *TDH* and *ctxA* detection with AIOD-CRISPR and PECAN, The AIOD-CRISPR utilises strand displacement to produce ssDNA, and the binding sites are exposed earlier if the target sequence is located near the primers that amplify the target strand. Therefore, the *TDH*1 crRNA design is advantageous to the AIOD-CRISPR system since the target site is close to the 5’ end of the amplicon. Upon binding of the phosphorothioate-protected primers, the strand displaced immediately provides a binding site for *TDH*1 crRNA. While for the *ctxA* target, the *ctxA*_R2 crRNA binds to the 3’ of the target strand of the amplicon, which delays the binding, and the ssDNA generated is more likely to be occupied by the opposite primer, decreasing the time available for crRNA binding, lowering the total activated cas12a. Furthermore, since the design of crRNAs in AIOD-CRISPR requires the crRNA binding site being adjacent to RPA primers. The design might limit the usage of AIOD-CRISPR based diagnosis due to the requirement of a consecutive 50bp conserved region for an efficient detection assay. However, as demonstrated, the PECAN system can utilise the whole amplicon efficiently for a flexible crRNA location design, as T7exo fully convert the dsDNA amplicon into ssDNA within a suitable condition (Supplementary Fig. 12).

## Discussion

Compared to the previous study utilising ssDNA as a cas12a activator without PAM activation [13], the cas12a concentration of the AIOD-CRISPR system is 1.28μM due to buffer incompatibility of cas12a in RPA buffer, while for PECAN it is 0.2μM. Previous research investigating the specificity of cas12a detection of ssDNA template suggested that with higher cas12a enzyme in the reaction, the less specific the cas12a trans-cleavage activity is, as high concentrations of cas12a were shown to decrease the minimum activating bases for crRNA complementary sequence [35]. The high cas12a usage also leads to a detection cost per reaction of $5.4 USD. The high detection cost decreases the desire for commercialisation and applicability of point of care (POC) detection at restaurants and farms. With a 6-fold reduction in cost for PECAN to $0.8 USD per reaction, the technology might be more economic and applicable.

While there are other strategies to generate ssDNA *in vitro* ssDNA for cas12a PAM-less activation, for example, the asymmetric PCR (aPCR) [43] and LAMP [44] assay could be utilised for ssDNA production. However, they require high reaction temperature and a thermal cycler for the operation, which limits the application outside of laboratories. Other strategies, including primer exchange [45] and rolling circle amplification [46], require a deep understanding of the assay, accompanied by complicated primer design. We also explored the strategy of asymmetric RPA (aRPA) [47] to generate ssDNA for cas12a activation. While the strategy has some success in producing high fluorescence signal compared to normal RPA reaction for cas12a PAM-less activation, we found that the result between different templates might not be consistent, and aRPA generated an unspecific amplification with the genomic template possibly due to a high concentration of primer, rendering the detection method less reliable (Supplementary Fig. 16). Therefore, we choose T7exo as the strategy for generating ssDNA templates for cas12a PAM-less activation due to its simplicity in design, with slight modifications of conventional RPA-cas12a detection methods, which is to include the T7exo enzyme and to use a phosphorothioate protected primer.

While both dsDNA with PAM site and ssDNA without PAM site could be utilised as a template for cas12a-based detection, and the minimum activation length of 14-bp crRNA is required for activation of indiscriminate trans-cleavage for both ssDNA and dsDNA, care should be taken in the design of crRNA and template type to avoid decreased specificity. For dsDNA, the hybridisation of 5’ crRNA nucleotides is essential to initiate the whole crRNA being hybridised to trigger indiscriminate ssDNA cleavage [48]. However, it was shown that in ssDNA, a mismatch of the 5’ end of crRNA could still lead to cas12a activation, leading to a minimum activation base of 11bp only in GC-rich crRNA, which is 3-bp shorter than using dsDNA. The same research group proposed some strategies; for example, using AT-rich crRNA target strand increases the specificity of ssDNA crRNA detection up to 14-bp minimum activation base [35]. This also suggests that the crRNA ssDNA hybridisation mechanism might be simple, like with the use of RPA fluorophore probe, which generates a free fluorophore per hybridised strand. In contrast, cas12a trans-cleavage could lead to a signal amplification per hybridised crRNA. Other strategies were also investigated, to increase the applicability of PECAN system to reliably generate fluorescence in the presence of the correct template. For crRNA engineering, a research group created the ENHANCE system, leading to increased specificity in case of SNP mutation. Moreover, ENHANCE [49] was also proven to work with ssDNA templates in our study to provide a >5-fold trans-cleavage activity with the addition of 3’7DNA to crRNA *TDH*1 without increased background (Supplementary Fig. 1), to achieve a brighter visual signal for cas12a trans-cleavage. Yet the Addition of 7-nt of DNA might hybridise with target ssDNA strand, and lead to concern to specificity. However, it was shown that with a crRNA extension of 5-nt, the minimum activating bases increased from 11-nt to 16-nt [35], indicating that 3’ extensions of crRNA might not decrease the required bases of activation of cas12a. The use of cas12a could therefore act as a secondary specificity safeguard, together with the RPA assay which is itself highly specific.

With PECAN, pathogens with small regions of conserved regions could be detected with cas12a without the presence of a conserved PAM site, which decreases the need for degenerative or polymorphic crRNA [50]. PECAN is applicable to the detection of bacterial or viral species. And illustrated in Supplementary Table 6, Norovirus GI only has a PAM-less 24-bp conserved region from sequence alignment for crRNA design, and another less conserved region for RPA primer design, which is not adjacent to the crRNA site. In our current research, we demonstrated the use of PECAN for easy-to-design, cheap and rapid DNA detection and demonstrated its adaptation for RT-RPA and RT-PCR for RNA detection.

## Methods

### Cloning

G-block containing a partial fragment of *TDH*, vcgC and *ctxA* (Supplementary Table 4a) was purchased and synthesised from IDT, and the gene block was cloned into XbaI and SpeI linearised pSB1C3 with NEBuilder (New England Biolab Inc.). PAM addition to *TDH* was carried out with mutagenesis primers (Supplementary Table 4C), and the PCR product from the primers was assembled with NEBuilder (New England Biolab Inc.). Plasmids with PAM site were screened with cas12a trans-cleavage assay without T7exo (Supplementary Fig. 13).

### *In vitro* transcription

*In vitro* transcription was carried out with MEGAshortscript™ T7 Transcription Kit (Invitrogen) according to the manufacturer’s guidelines. For crRNA transcription, 100μM of crRNA template strand with T7 promoter binding site were mixed with 100μM of T7_anneal (Supplementary Table 2a) in 2x NEB r2.1 buffer. The mixture was annealed in a thermocycler with a gradual decrease in temperature over 2 hours. The annealed product (40μM) was diluted with water to a final concentration of 10μM and added to the T7 MEGAshortscript transcription mix to a final concentration of 1μM. The mixture was then incubated at 37 degrees for 24 hours. For the transcription of the *NoV GI* synthetic RNA template, the *NoV GI* DNA template was synthesised from IDT with T7 promoter and was added to T7 MEGAshortscript transcription complete reaction to a final concentration of 0.1μM and incubated for 24 hours. After the transcription reaction, Turbo DNase was added to the transcription reaction and incubated for 1 hour. The transcription reaction was either purified by MEGAclear Transcription Clean-Up Kit (Invitrogen) or higher yield with phenol-chloroform extraction followed by ethanol precipitation according to the manufacturer’s instruction. The crRNA is eluted in TE buffer and frozen for storage, and the concentration was measured by nanodrop and adjusted to 20μM. The crRNA quality was analysed with 15% SDS-UREA PAGE with a 19:1 ratio for cross-linker ran at room temperature with a low current. The crRNA was stained with 1x SYBR gold solution until RNA was visualised under blue light or UV transilluminator (Supplementary Fig.15).

### Design of crRNA and primers

FASTA files for gene targets were downloaded from the NCBI Nucleotide database with search terms outlined in Supplementary Table 6. The FASTA files were trimmed away from unrelated sequences with overly short sequences, promoter region, genome shotgun and partial cds. Then the FASTA files were uploaded to Cluster Omega for sequence alignment with ClusterW with character count. The output alignment was visualised with Jalview to identify conserved nucleotides. crRNA candidates were designed within sequences with complete consensus while flanking by RPA primers in a conserved site, ideally with amplicon size between 100-200bp. The crRNA candidates were screened with CRISPR-DT [51] to obtain a crRNA score and the highest score candidate. However, we also screened crRNA without CRISPR-DT by selecting a 20-nt sequence without a loss of activity of crRNA. For RPA primers, we followed instructions from TwistDx and designed primers with GC content of 40-60%, with a length between 30-32 nt. The RPA primers cross, and self-hybridisation were checked with a pre-setting of 3 on the Multiple Primer Analyser webtool of ThermoFisher. The secondary structure of RPA primers and crRNA was checked with IDT OligoAnalyzer Tool. Finally, the shortlisted candidates were input into BLAST to identify potential false positives of selected crRNA.

### PECAN assay

5x PECAN “Mix” reaction buffer was assembled with 25μL of NEB r2.1 (10x), 25μL of NEB 4 (10x), and 50μL of 10mM DTT per 100μL of 5x buffer. EnGen Lba Cas12a 100μM were purchased from New England Biolab Inc. and were diluted into 10μM stock with the diluent provided with the enzyme for small-scale experiments. For cas12a-crRNA coupling, 10μM cas12a was mixed 1:1 with 20μM crRNA and incubated at least 10-min at room temperature before use. After the 10-min incubation, the PECAN reaction mix was assembled as described. Per 10μL of PECAN reaction, 0.1μL (1 Unit) T7exo, and 0.4μL (final cas12a concentration of 0.2μM) of cas12a-crRNA mix, 0.8μL of 100μM ssDNA-FQ, and a final of 1X Mix Buffer were added. RPA reaction was carried out with TwistAmp Liquid Basic Kit (TwistDx Limited) with 0.32μM of each primer and 1.8mM of dNTP. For RPA PECAN, 0.5μL of the 20-min RPA reaction was directly added to 10μL PECAN without purification. Then the PECAN reaction was incubated at 37 degrees for 20-min for fluorescence visualisation. After incubation, 1μL of 0.5M EDTA solution was added to each tube to quench and stop the reaction. The fluorescence endpoint was measured in a 384-well black plate with an optical bottom (Greiner) with Cytation 1 (BioTek), or visualized with a blue light or UV transilluminator. For gel analysis of T7exo activity in PECAN and visualisation of dsDNA/ssDNA produced by RPA or RPA-PECAN, the assay mixture was loaded into a 3% agarose gel with 1x SYBR Gold (Invitrogen) and visualised with a blue light or UV transilluminator.

## Acknowledgements

The iGEM impact grant for team Hong Kong International School (HKIS) funded the study.

## Author contributions

K.N. performed the bioinformatics analysis of the gene targets and designed the nucleic acid sequences. All authors performed pilot test for RPA, asymmetric RPA, cas12a, T7exo and DNA gel analysis. K.N., D.Z. performed and optimised the PECAN detection reactions, analysed the data. D.Z. and S.R. conducted the genomic DNA RPA and lateral flow assay. K.N. made the figures. All authors edited the manuscript.

