## Supplementary Information and figures for "PAM-less Exonuclease-assisted Cas12a for visual detection of Vibrio Species"

### 2Supplementary Information

**Supplementary Table 1: Sequences of crRNA used in this study.** The crRNA and the direction relative to the RPA/PCR primers are denoted (e.g., reverse direction of crRNA is complementary to the strand generated by forward primer). *Denote the crRNA used for the final PECAN detection. The last 20nt (red) are the complementary sequences of the target sequence, while before that is the cas12a crRNA stem-loop. Note that with T7 transcription 3-extra G residues would be present at the 5' end.

| crRNA name | Direction | PAM | Sequence |
|---|---|---|---|
| TDH1 * | Forward | - | GGGUAAUUUCUACUAAGUGUAGAUACUGUGAACAUUAAUGAUAA |
| TDH_R | Reverse | - | GGGUAAUUUCUACUAAGUGUAGAUGUUGUGAAUACUGAUUGACC |
| TDH2 * | Forward | - | GGGUAAUUUCUACUAAGUGUAGAUGGUCAAUCAGUAUUCACAAC |
| vcgC_F * | Forward | - | GGGUAAUUUCUACUAAGUGUAGAUAUAGCACUAAUGUGUCAUCU |
| vcgC_R | Reverse | + | GGGUAAUUUCUACUAAGUGUAGAUGCGGAGACGAGAUCGCUAUC |
| ctxA_F1 | Forward | + | GGGUAAUUUCUACUAAGUGUAGAUUGAUCAUGCAAGAGGAACUC |
| ctxA_R1 | Reverse | + | GGGUAAUUUCUACUAAGUGUAGAUAGUACCUCGGUCAAAGUACU |
| ctxA_R2 * | Reverse | - | GGGUAAUUUCUACUAAGUGUAGAUGAGUUCCUCUUGCAUGAUCA |
| ctxA_R3 | Reverse | - | GGGUAAUUUCUACUAAGUGUAGAUAUUCAUUUGAGUACCUCGGU |
| NoVGI_R | Reverse | - | GGGUAAUUUCUACUAAGUGUAGAUCCUUAGACGCCAUCAUCAUU |
| NoVGI_F * | Forward | - | GGGUAAUUUCUACUAAGUGUAGAUGAUGAUGGCGUCUAAGGACG |
| TDH1_ 3'7DNA * | Forward | - | rUrArArUrUrUrCrUrArCrUrArArGrUrGrUrArGrArUrArCrUrGrUrGrArArCrArUrUrArArUrGrArUrArATATTATT |

The final crRNA TDH1_3'7DNA is a crRNA/DNA version of TDH_1 with a 3' 7 nt DNA extension with ENHANCE system [1], providing increased trans-cleavage activity. ctxA_R2 showed no trans-cleavage activity towards ssDNA template.

**Supplementary Table 2: Sequences of template for transcription of crRNA used in this study.** (a) The crRNA and the direction relative to the RPA/PCR primers are denoted (reverse direction of crRNA is complementary to the strand generated by forward primer). The template is reverse complement to the crRNA, with a binding site for a T7 promoter such that the annealed product could be utilized for *in vitro* transcription from DNA template to crRNA. First 20nt (Red) = crRNA complementary sequence. Last 25nt (Blue) = T7 promoter annealing site. (b) Illustration of annealing and the direction of transcription. The transcript would start at the T7 promoter with GGG, extending to the stem-loop and to the crRNA complementary region, with direction the same as T7_anneal.

## a

| Target | Direction | PAM | Sequence |
|---|---|---|---|
| TDH_1 | Forward | - | TTATCATTAATGTTCACAGTATCTACACTTAGTAGAAATTACCC TATAGTGAGTCGTATTAATTTC |
| TDH_R | Reverse | - | GGTCAATCAGTATTCACAACATCTACACTTAGTAGAAATTACCC TATAGTGAGTCGTATTAATTTC |
| TDH_2 | Forward | - | GTTGTGAATACTGATTGACCATCTACACTTAGTAGAAATTACCC TATAGTGAGTCGTATTAATTTC |
| vcgC_F | Forward | - | AGATGACACATTAGTGCTATATCTACACTTAGTAGAAATTACCC TATAGTGAGTCGTATTAATTTC |
| vcgC_R | Reverse | + | GATAGCGATCTCGTCTCCGCATCTACACTTAGTAGAAATTACCC TATAGTGAGTCGTATTAATTTC |
| ctxA_F1 | Forward | + | GAGTTCCTCTTGCATGATCAATCTACACTTAGTAGAAATTACCC TATAGTGAGTCGTATTAATTTC |
| ctxA_R1 | Reverse | + | AGTACTTTGACCGAGGTACTATCTACACTTAGTAGAAATTACCC TATAGTGAGTCGTATTAATTTC |
| ctxA_R2 | Reverse | - | TGATCATGCAAGAGGAACTCATCTACACTTAGTAGAAATTACCC TATAGTGAGTCGTATTAATTTC |
| ctxA_R3 | Reverse | - | ACCGAGGTACTCAAATGAATATCTACACTTAGTAGAAATTACCC TATAGTGAGTCGTATTAATTTC |
| NoVGI_R | Reverse | - | AATGATGATGGCGTCTAAGGATCTACACTTAGTAGAAATTACCC TATAGTGAGTCGTATTAATTTC |
| NoVGI_F | Forward | - | CGTCCTTAGACGCCATCATCATCTACACTTAGTAGAAATTACCC TATAGTGAGTCGTATTAATTTC |
| T7_anneal | | | GAAATTAATACGACTCACTATAGGG |

## b

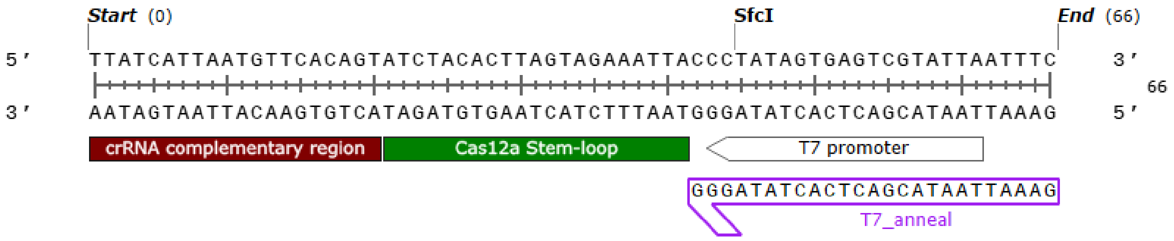

**Supplementary Table 3: Sequences of primers used in this study.** (a) RPA primers designed for all the gene targets. (b) qPCR primers used for *Vibrio species*.

## a

| Gene target | Organism | Direction | Sequence |
|---|---|---|---|
| TDH | *Vibrio parahaemnolyticus* | Forward | CCCGGTTCTGATGAGATATTGTTTGTTGTTCG |
| TDH | *Vibrio parahaemnolyticus* | Reverse | CTTATAGCCAGACACCGCTGCCATTGTATAGT |
| vcgC | *Vibrio Vulnificus* | Forward | GGCGCAGTTCAAACATGGTCTCAAAAAGGAGC |
| vcgC | *Vibrio Vulnificus* | Reverse | GCGAGTAGTGAGCCGATCCCGCCAATAGCCTG |
| ctxA | *Vibrio Cholerae* | Forward | GCAGTCAGGTGGTCTTATGCCAAGAGGACAGA |
| ctxA | *Vibrio Cholerae* | Reverse | ACATATCCATCATCGTGCCTAACAAATCCCGT |
| NoVGI | *Norovirus GI* | Forward | TGGCAGGCCATGTTTCGCTGGATGCG |
| NoVGI | *Norovirus GI* | Reverse | GGTCAGAAGTATTAGCCTCCGGTACCA |
| NoVGI_Degen | *Norovirus GI* | Reverse | TCAGMWGTATTWRCCTCYGGTACCA |

## b

| | |
|---|---|
| tdh_qPCR_F | CCCGGTTCTGATGAGATATTGT |
| tdh_qPCR_R | GACACCGCTGCCATTGTATAGT |
| vcgC_qPCR_F | GCAGTTCAAACATGGTCTCAAA |
| vcgC_qPCR_R | CGCCAATAGCCTGTTCAGAT |
| ctxA_qPCR_F | GGTCTTATGCCAAGAGGACAGA |
| ctxA_qPCR_R | CCATCATCGTGCCTAACAAATC |

**Supplementary Table 4: Sequences of primers used in this study.** (a) DNA templates of partial sequences of *Vibrio* species pathogenic marker *TDH* (First highlight, yellow), *vcgC* (Second highlight, green) and *ctxA* (Third highlight, blue) to be cloned into pSB1C3 as a biobrick. And the DNA template for in vitro transcription of partial genome of *norovirus I* with a T7 promoter at the 5' end. (b) T7 promoter transcribed NoV GI genomic RNA mimic. (c) Primers for mutation and adding a TTTA PAM site before TDH1 crRNA for a PAM site positive control. (d) Insert after mutagenesis (PAM site highlighted in reverse direction).

**a**

| | |
|---|---|
| Vibrio_TDH_vcgC_ctxA | gaattcgcggccgcttctagagCTTATAGCCAGACACCGCTGCCATTGTATAGTCTTTATCATTAATGTTCACAGTCATGTAGGATGTCAACCATTTAGTACCTGACGTTGTGAATACTGATTGACCATAAACATCTTCGTACGGTTTTCTTTTTACATTACGGTTTGTCCAAAAGTCAGAGACCTTTACATTGACCGGAGCTTGGGTATTAAAAGTTGTATCTCGAACAACAAACAATATCTCATCAGAACCGGGagatctgtcgacGCGAGTAGTGAGCCGATCCCGCCAATAGCCTGTTCAGATGACACATTAGTGCTATTTTGTATCGCGGAGACGAGATCGCTATCGGCAGCTCCTTTTTGAGACCATGTTTGAACTGCGCCaagcttGCAGTCAGGTGGTCTTATGCCAAGAGGACAGAGTGAGTACTTTGACCGAGGTACTCAAATGAATATCAACCTTTATGATCATGCAAGAGGAACTCAGACGGGATTTGTTAGGCACGATGATGGATATGTGCTAGCtgtacaggatcctactagtagcggccgctgcag |
| T7_NoVGI | GAAATTAATACGACTCACTATAGGGAgaCCAGCAAAGTCATACATGAAATCAAGACTGGTGGATTGGAGATGTATGTTCCAGGGTGGCAGGCCATGTTTCGCTGGATGCGCTTCCATGACCTCGGATTGTGGACAGGAGATCGCAATCTCCTGCCCGATTTCGTAAATGATGATGGCGTCTAAGGACGCTACGTCAAACGTGGATGGCGCCAGCGGCGCTGGTCAGTTGGTACCGGAGGCTAATACTTCTGACCCCCTTGCAATGGATCCTGTGGCGGGTTCTTCGACAGCGGTTGC |

Reference accession: TDH: CP020428.2 , vcgC: CP014049.2, ctxA: CP047059, NoV GI: MT008453.1

**b**

| | |
|---|---|
| RNA_NoVGI | GGGAGACCAGCAAAGUCAUACAUGAAAUCAAGACUGGUGGAUUGGAGAUGUAUGUUCCAGGGUGGCAGGCCAUGUUUCGCUGGAUGCGCUUCCAUGACCUCGGAUUGUGGACAGGAGAUCGCAAUCUCCUGCCCGAUUUCGUAAAUGAUGAUGGCGUCUAAGGACGCUACGUCAAACGUGGAUGGCGCCAGCGGCGCUGGUCAGUUGGUACCGGAGGCUAAUACUUCUGACCCCCUUGCAAUGGAUCCUGUGGCGGGUUCUUCGACAGCGGUUGC |

**c**

| | |
|---|---|
| VP_TDH_mut_F | TCACAGTtaaaTAGGATGTCAACCATTTAGTACCTGAC |
| VP_TDH_mut_R | CATCCTAtttaACTGTGAACATTAATGATAAAGACTATACAATGGC |

**d**

| | |
|---|---|
| Vibrio_PAM_TDH | gaattcgcggccgcttctagagCTTATAGCCAGACACCGCTGCCATTGTATAGTCTTTATCATTAATGTTCACAGTtaaaTAGGATGTCAACCATTTAGTACCTGACGTTGTGAATACTGATTGACCATAAACATCTTCGTACGGTTTTCTTTTTACATTACGGTTTGTCCAAAAGTCAGAGACCTTTACATTGACCGGAGCTTGGGTATTAAAAGTTGTATCTCGAACAACAAACAATATCTCATCAGAACCGGGagatctgtcgacGCGAGTAGTGAGCCGATCCCGCCAATAGCCTGTTCAGATGACACATTAGTGCTATTTTGTATCGCGGAGACGAGATCGCTATCGGCAGCTCCTTTTTGAGACCATGTTTGAACTGCGCCaagcttGCAGTCAGGTGGTCTTATGCCAAGAGGACAGAGTGAGTACTTTGACCGAGGTACTCAAATGAATATCAACCTTTATGATCATGCAAGAGGAACTCAGACGGGATTTGTTAGGCACGATGATGGATATGTGCTAGCtgtacaggatcctactagtagcggccgctgcag |

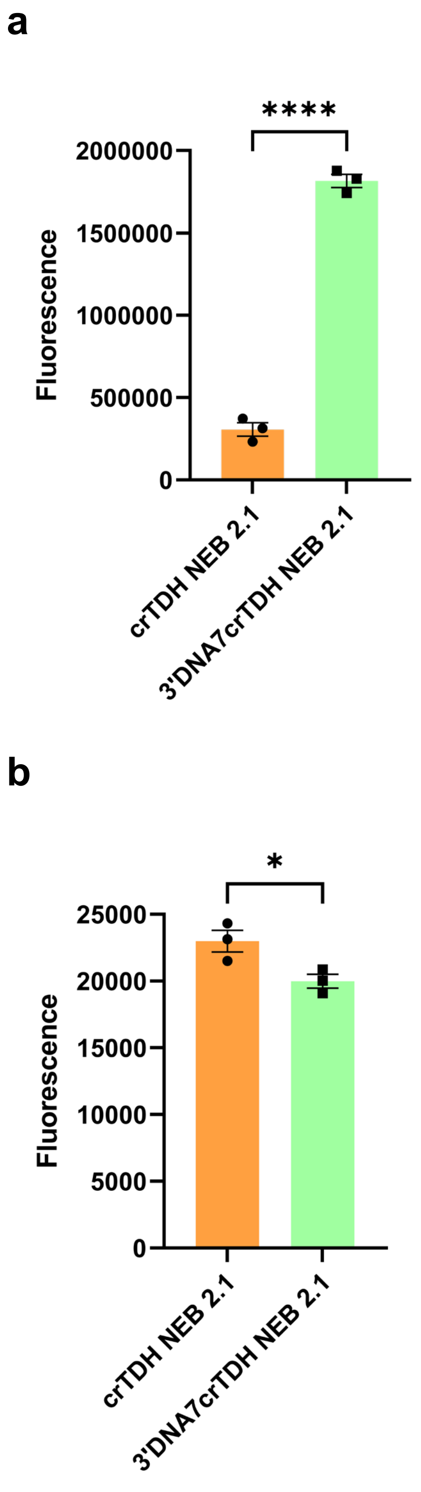


**Supplementary Figure 1. a** Trans-cleavage activity of cas12a with crRNA TDH1, and with crRNA with 3'7DNA modification. **b** And the background cleavage activity. The unpaired two-tailed t-test was used to analyze the statistical significance.

**a**

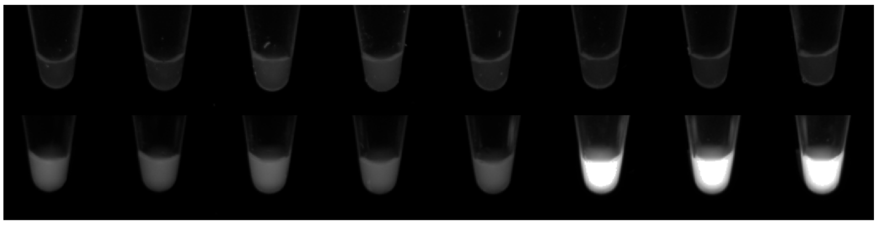


| Buffer | NEB2.1 | Mix | Mix + 1mM DTT | Mix + 10mM DTT | Mix + 1mM TCEP | NEB2.1 | Mix | Mix + 1mM DTT |
|---|---|---|---|---|---|---|---|---|
| DTT conc. | 0 | 0.5 | 1.5 | 10.5 | | 0 | 0.5 | 1.5 |
| crRNA | crTDH | crTDH | crTDH | crTDH | crTDH | crTDH+ 3'7DNA | crTDH+ 3'7DNA | crTDH+ 3'7DNA |

**b**

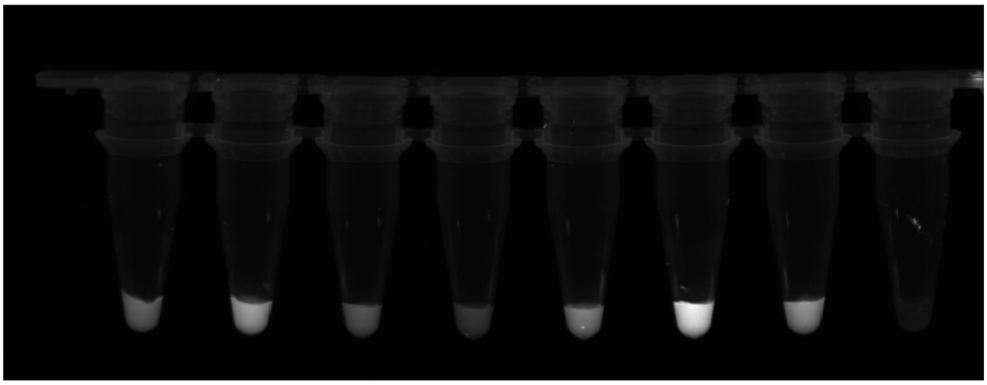

| cas12a | + | + | + | + | + | + | + | - |
|---|---|---|---|---|---|---|---|---|
| Buffer composition | r2.1 (1x) | r2.1 (0.5x) + 4 (0.5x) + 0.5mM DTT | r2.1 (1x) + 4 (1x) | 4 (1x) | 4 (1x) + 50mM NaCl | 4 (1x) + 10mM MgCl2 | 4 (1x) | r2.1 (1x) |

**Supplementary Figure 2. a** Trans-cleavage activity of cas12a with crRNA TDH1, and with crRNA with 3'7DNA modification. **b** Trans-cleavage activity of cas12a in various buffer. 0.1μM of cas12a were added to 1μM of ssDNA template with 8μM of ssDNA-FQ reporter.

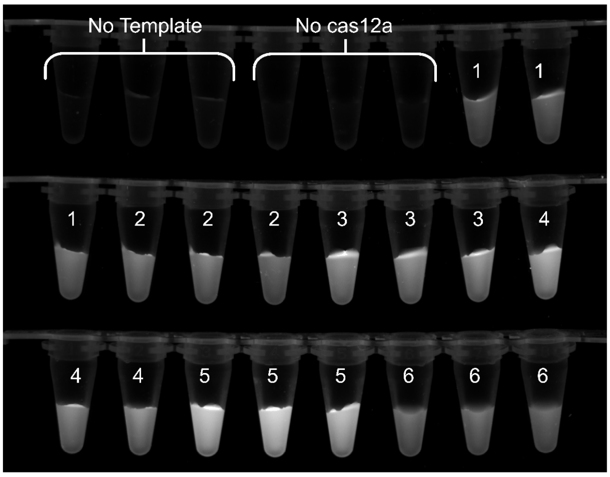


| Number | 1 | 2 | 3 | 4 | 5 | 6 |
|---|---|---|---|---|---|---|
| cas12a | + | + | + | + | + | + |
| Buffer composition | r2.1 (1x) | r2.1 (0.5x) + 4 (0.5x) | r2.1 (0.5x) + 4 (0.5x) | r2.1 (0.5x) + 4 (0.5x) | r2.1 (0.5x) + 4 (0.5x) | 4 (1x) |
| DTT (mM) | 0 | 0.5 | 1 | 0.5 | 1 | 1 |
| Mg2+ | 10 | 10 | 10 | 20 | 20 | 10 |

**Supplementary Figure 3.** Triplicate buffer analysis for cas12a trans-cleavage activity. 0.1µM of cas12a were incubated with 1µM of ssDNA target with 8µM of ssDNA-FQ as reporter for 20 minutes. The mixture was later diluted to 50µL for more accurate fluorescence visualization.

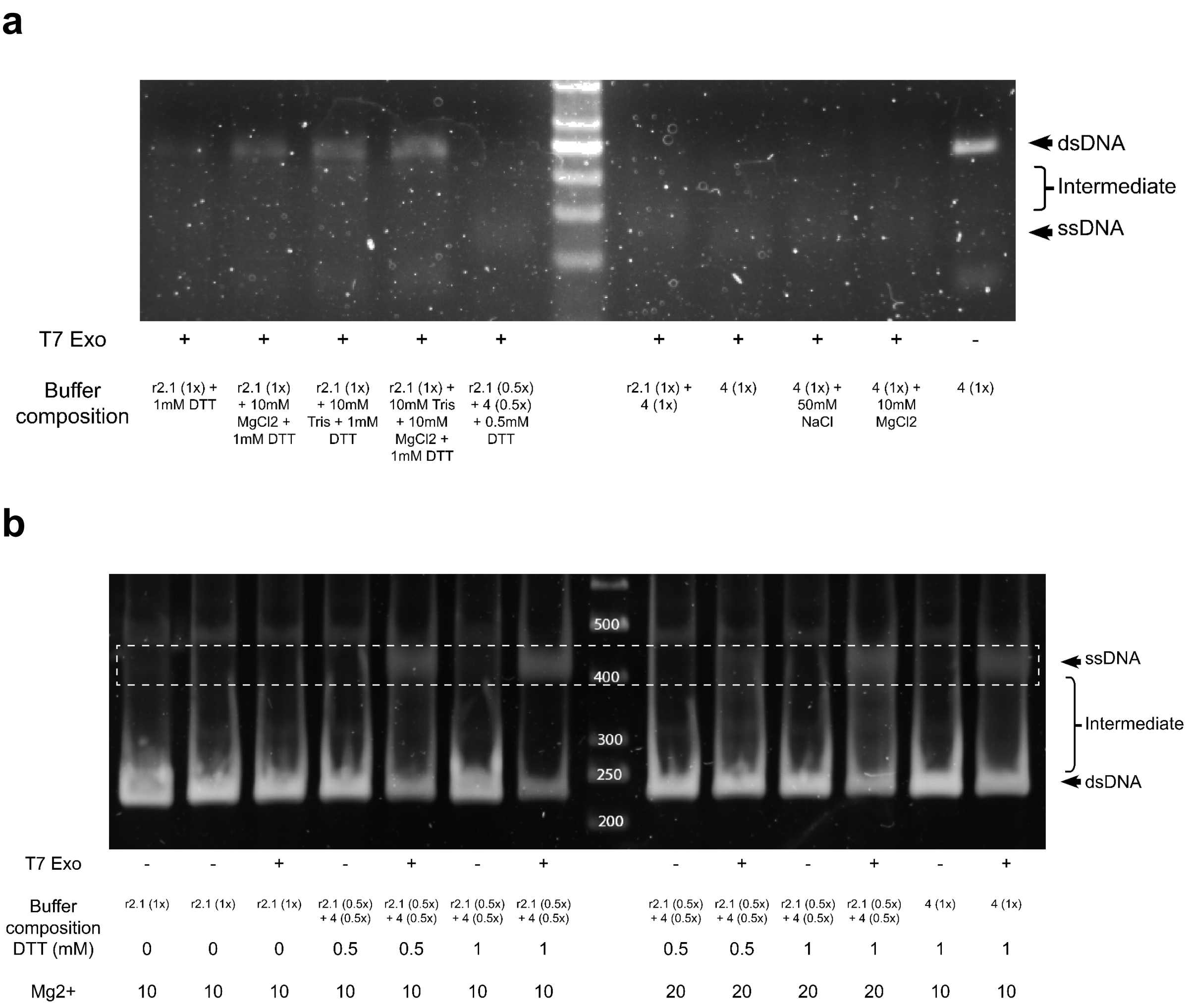


**Supplementary Figure 4. a** T7exo activity buffer analysis with 0.5µL of 20-min RPA in 10µL of reaction with 5 units of T7exo for 10-min. **b** T7exo activity buffer analysis with 100ng of PCR amplicon in 10µL of reaction with 1 units of T7exo in 20 min, visualized in 10% PAGE gel. Note that the banding of ssDNA and dsDNA shifted, due to the difference in pre-staining in agarose gel and post-staining in PAGE gel.

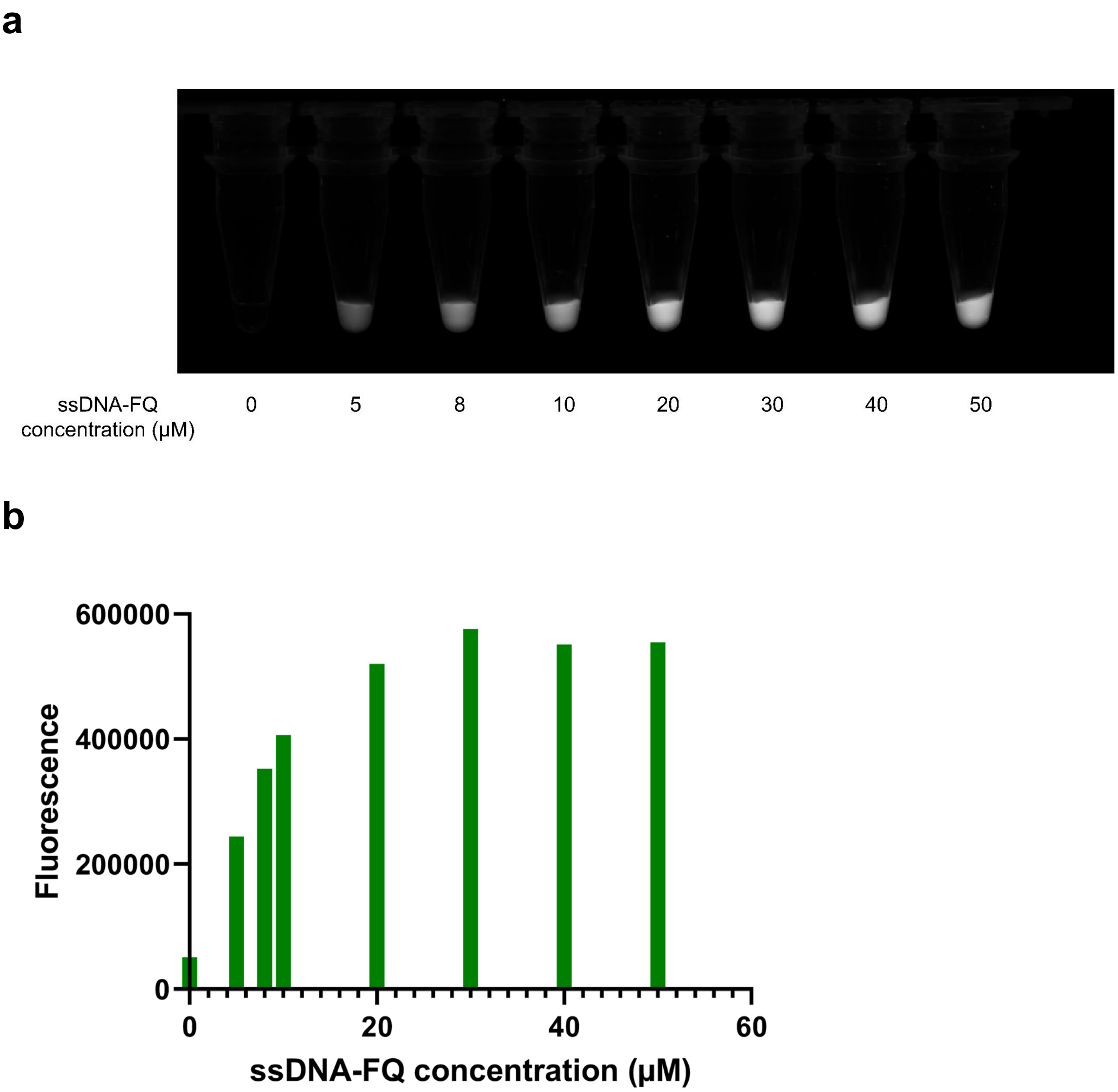


**Supplementary Figure 5.** ssDNA-FQ concentration analysis for cas12a trans-cleavage activity. The reaction was carried out in Mix buffer with 0.1μM of cas12a, 1μM of ssDNA template and varied amount of fluorophore. The reaction is carried out for 10-min at 37-degrees and quenched with the addition of EDTA. The analysis suggest that the linear relationship between ssDNA-FQ and fluorescence is between 0-10μM.

a

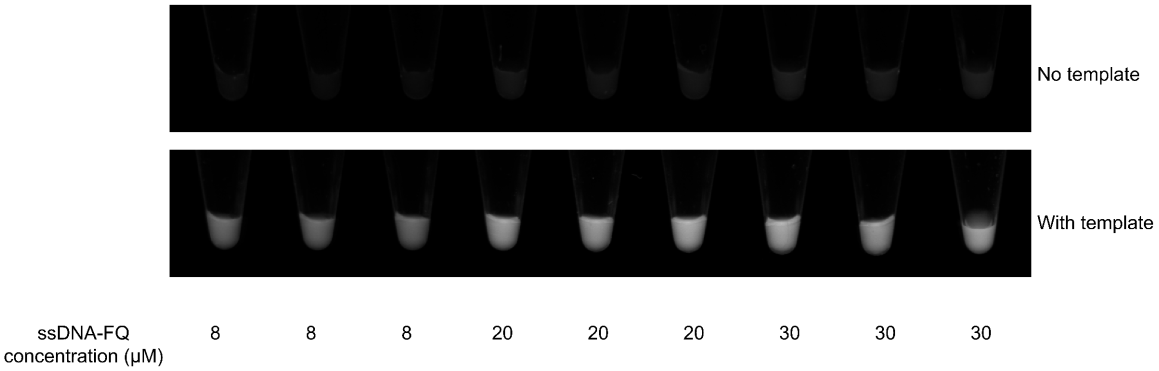


b

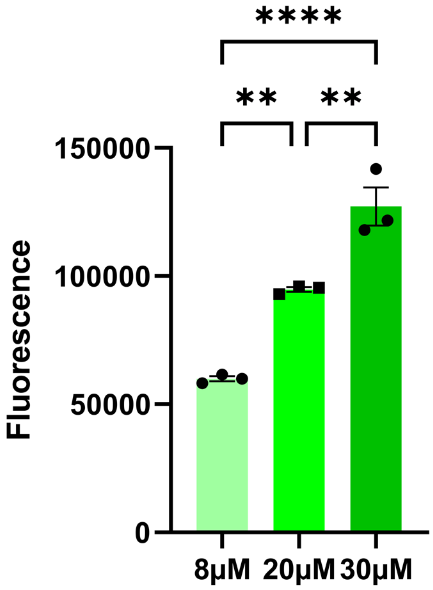


**Supplementary Figure 6. a** Triplicate trans-cleavage activity of cas12a with different fluorophore concentration with 0.1μM of cas12a in Mix buffer with 1μM of ssDNA as template. **b** And the background cleavage activity of different fluorophore concentration without template. Ordinary one-way ANOVA was used to analyze the statistical significance.

**a**

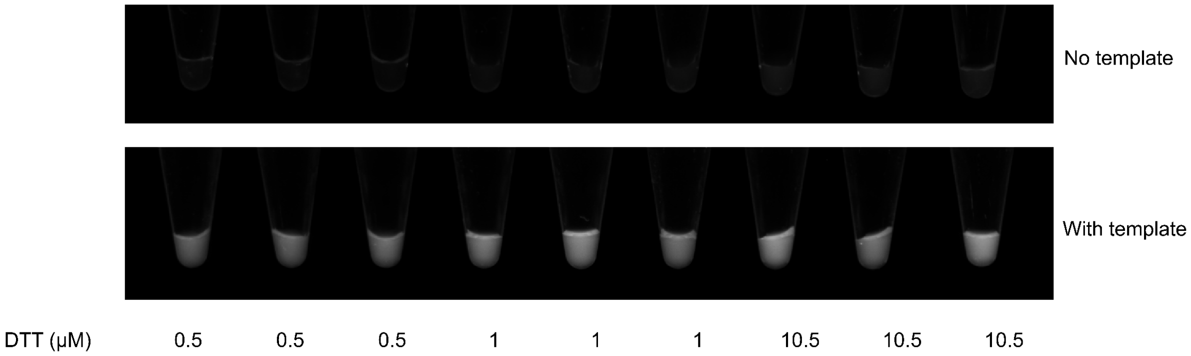


**b**

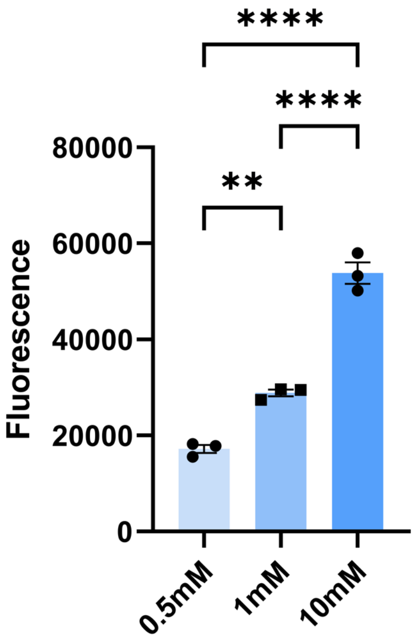


**Supplementary Figure 7. a** Triplicate trans-cleavage activity of cas12a with different DTT concentration with 0.1μM of cas12a in Mix buffer with 1μM of ssDNA as template. **b** And the background cleavage activity of different DTT concentration without template. Ordinary one-way ANOVA was used to analyze the statistical significance.

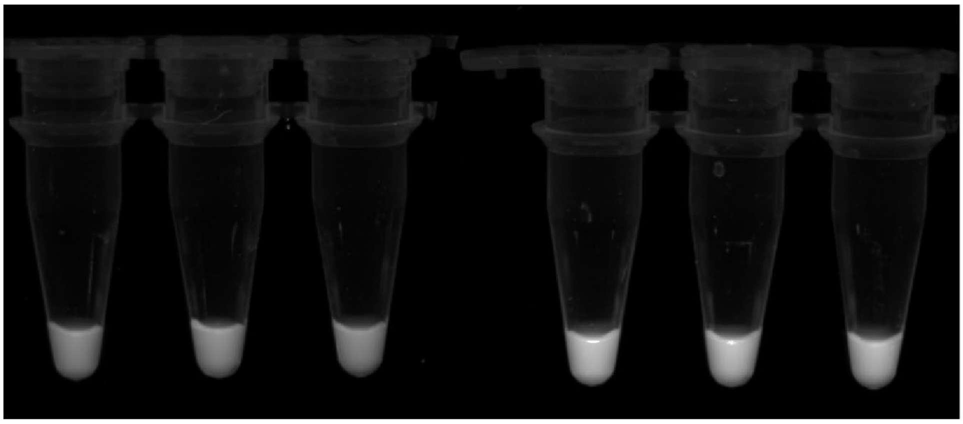

| | | | | | | |
|---|---|---|---|---|---|---|
| crRNA: cas12a | 1 | 1 | 1 | 2 | 2 | 2 |
| Incubation time (min) | 10 | 10 | 10 | 10 | 10 | 10 |

**Supplementary Figure 8.** Triplicate trans-cleavage activity of cas12a with different ratio of cas12a-crRNA 10-min coupling before added to the one pot reaction, with 0.1μM of cas12a in Mix buffer with 1μM of ssDNA as template.

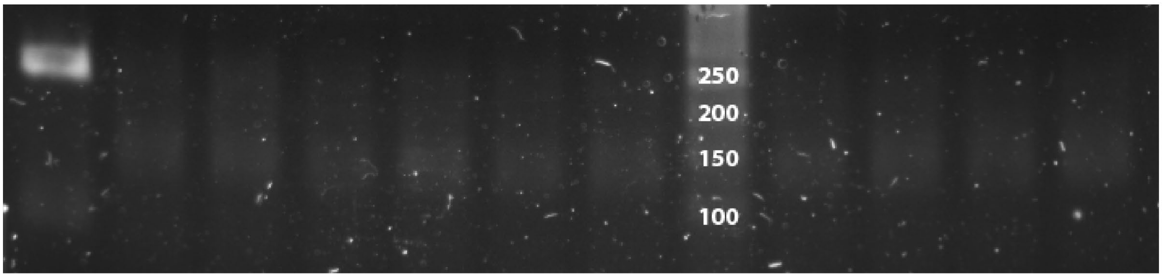


| | | | | | | | | | | | | |
|---|---|---|---|---|---|---|---|---|---|---|---|---|
| T7exo | 0 | 0.1 | 0.1 | 0.2 | 0.2 | 0.3 | 0.3 | / | 0.4 | 0.4 | 0.5 | 0.5 |
| Incubation time (min) | 20 | 10 | 20 | 10 | 20 | 10 | 20 | / | 10 | 20 | 10 | 20 |

**Supplementary Figure 9.** T7exo digestion gel analysis with 0.5μL of 20-min RPA added to 10μL of reaction with various units of T7exo for 10 or 20-min. 0.1μL indicates 1unit of T7exo was used.

**a**

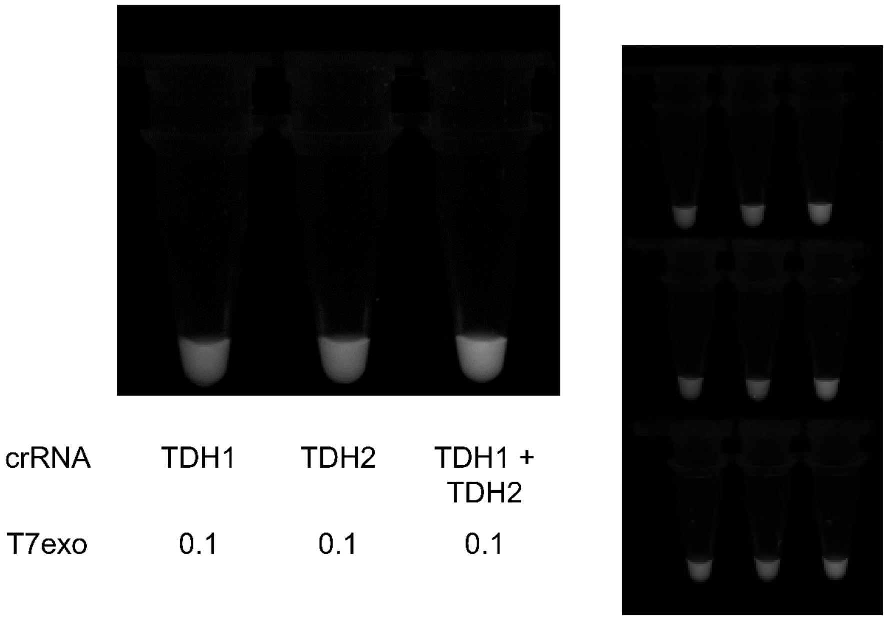


**b**

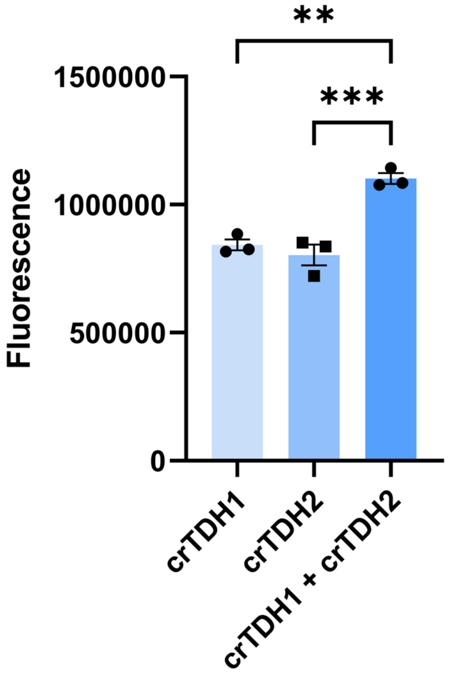


**Supplementary Figure 10. a** Triplicate trans-cleavage activity of cas12a with either single RNA or dual RNA in PECAN visualized in UV with 0.2µM of cas12a and 0.5µL of 20-min RPA amplificaition. **b** The fluorescence of (a). Ordinary one-way ANOVA was used to analyze the statistical significance.

**a**

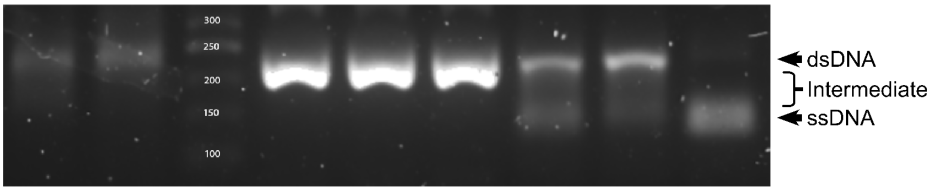


**b**

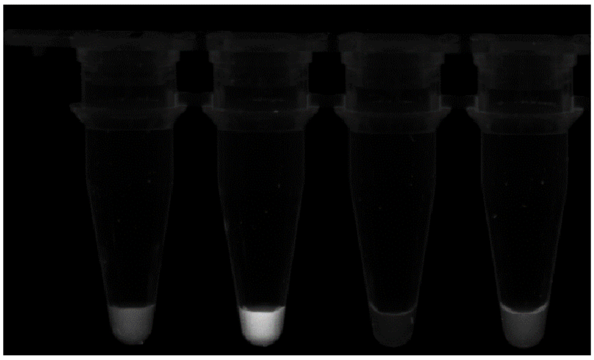


**Supplementary Figure 11. a** T7exo analysis with different 20-min heat-inactivated 20-min RPA reactions in different temperature. The ssDNA and dsDNA were denoted with arrows. 0.1µL (1Unit) of T7exo was added for each indicated reaction in a 10µL reaction, and incubated for 20-min in 37C before loading to agarose gel. **b** PECAN visualization and fluorescence produced with various 20-min heat-denatured 20-min RPA reactions.

**a**

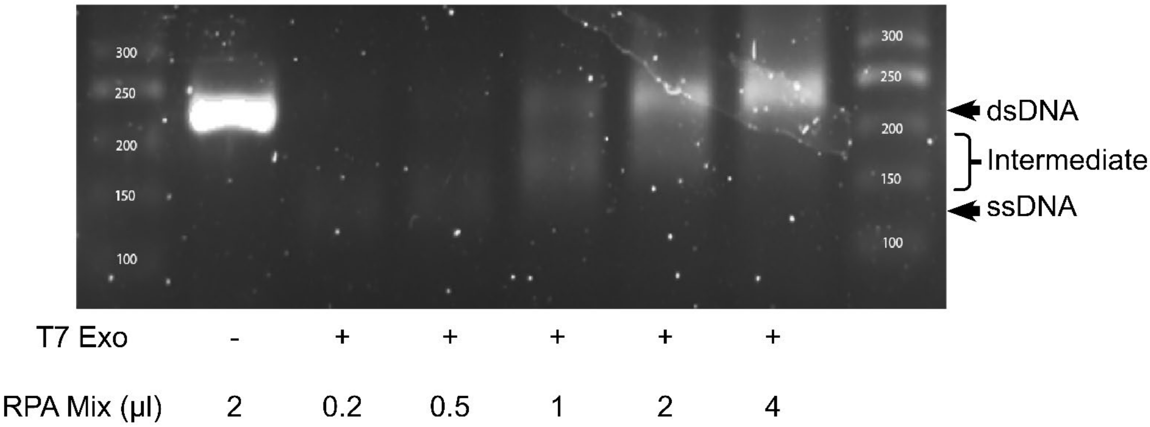


**b**

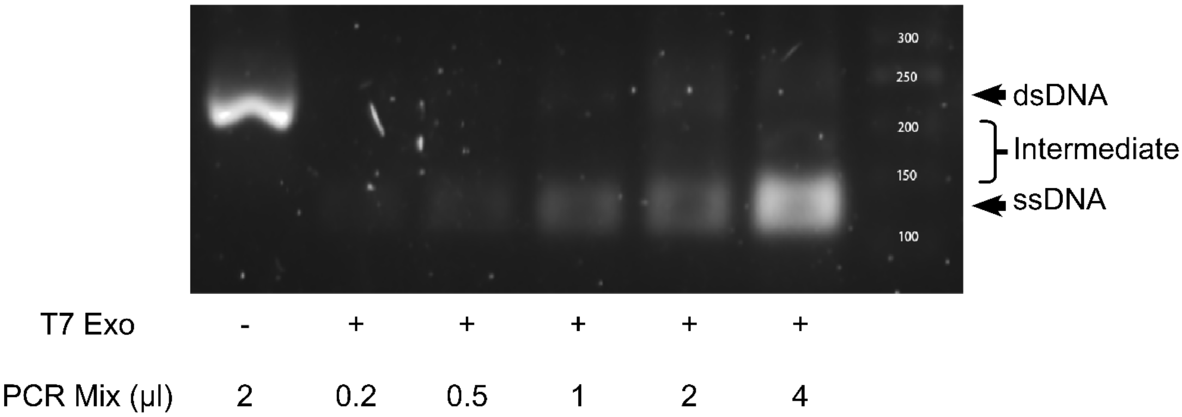


**Supplementary Figure 12. a** T7exo analysis with 20-min RPA reactions. Various volume of RPA reaction were digested with 0.1µL (1Unit) of T7exo in a 10µL reaction in Mix buffer, and incubated for 20-min in 37C before loading to agarose gel. **b** T7exo analysis with PCR endpoints. Various volume of PCR reaction were digested with 0.1µL (1Unit) of T7exo in a 10µL reaction in Mix buffer, and incubated for 20-min in 37C before loading to agarose gel.

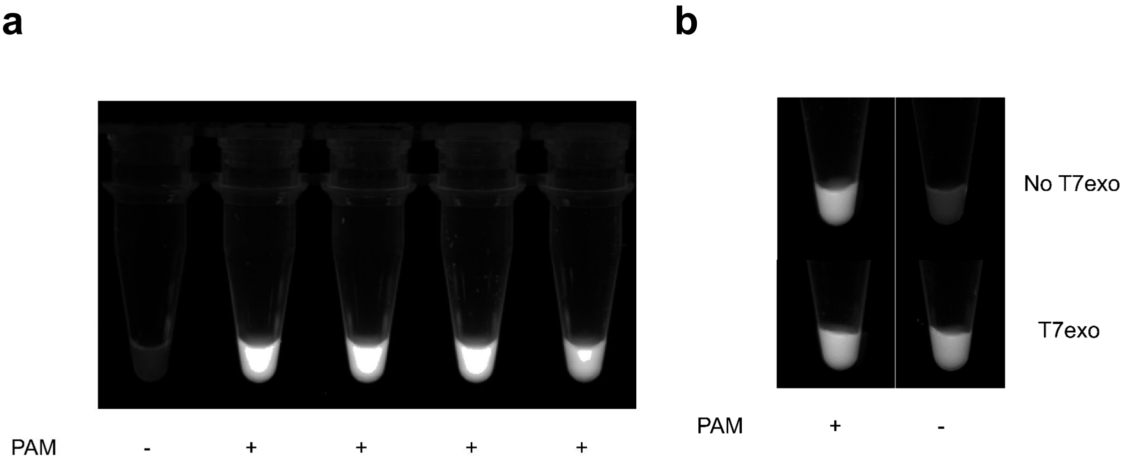


**Supplementary Figure 13. a** Cloning verification of addition of PAM site with site directed mutagenesis. 0.5μL of 20-min RPA of 4 miniprepped mutagenesis colonies were incubated with 0.1μM of cas12a with 8μM fluorophore without T7exo, while the 100ng of original plasmid was used as PAM-less control. **b** 1ng of plasmid was used as a template for RPA amplification and PECAN reaction with or without T7exo was used to analyse the fluorescence produced by PAM or PAM-less plasmid. The result indicates that without T7exo, PAMless plasmid would have a weak fluorescence, but it does not affect the plasmid with PAM. Indicating only when the PAM-less amplicon was transformed to ssDNA, it can produce a trans-cleavage activity.

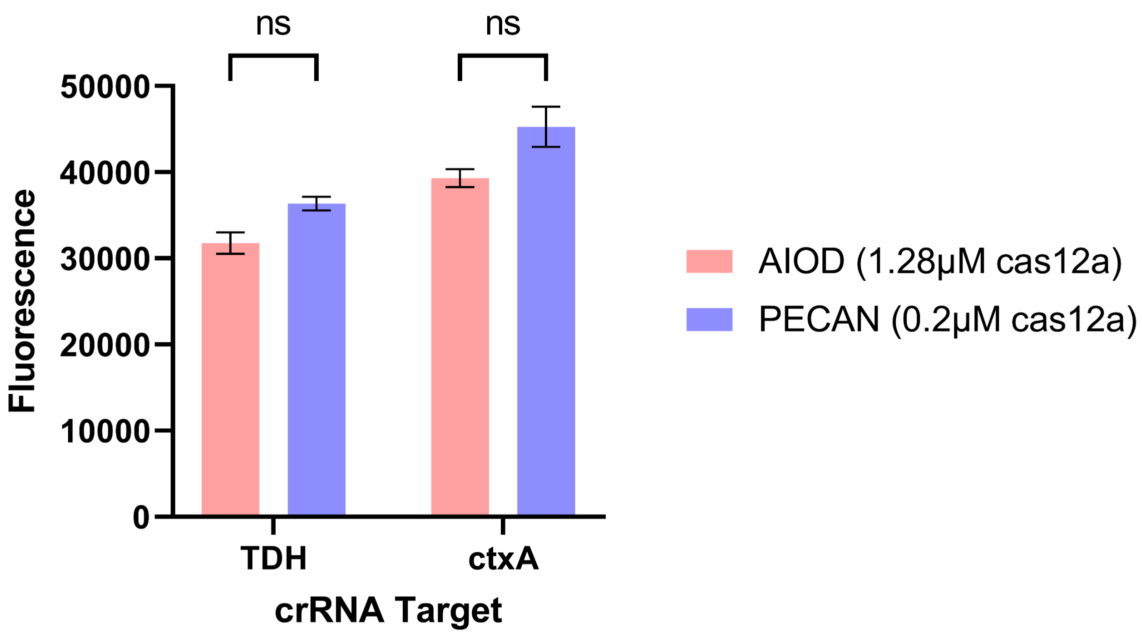


**Supplementary Figure 14.** Triplicate trans-cleavage activity of AIOD and PECAN in the absence of template (background signal). The result indicates that both system has negligible difference of background at the 40-min endpoint. Multiple two-tailed t-test was used to analyze the statistical significance.

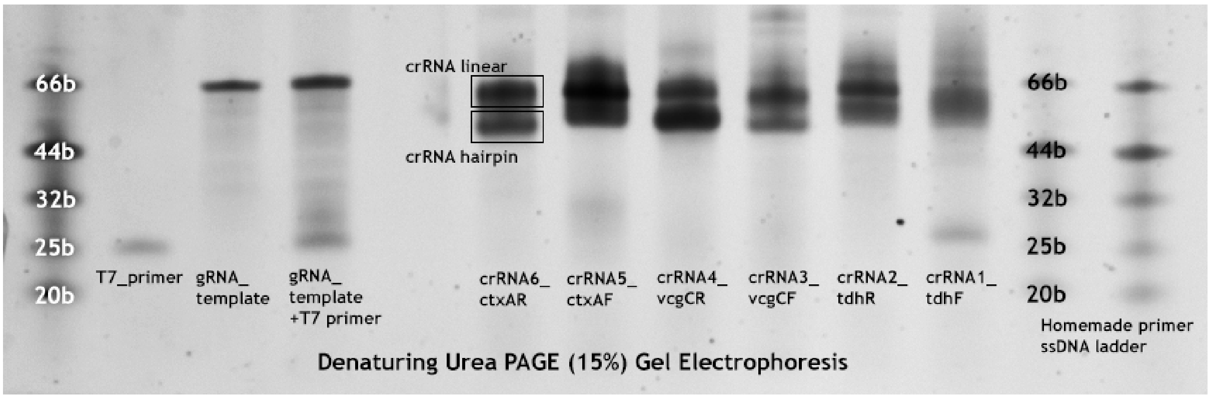


**Supplementary Figure 15.** 15% UREA PAGE gel analysis of crRNA transcription reaction with home-made ssDNA ladder made of DNA primers for relative base pair comparison. The gel was stained with SYBR gold.

**a**

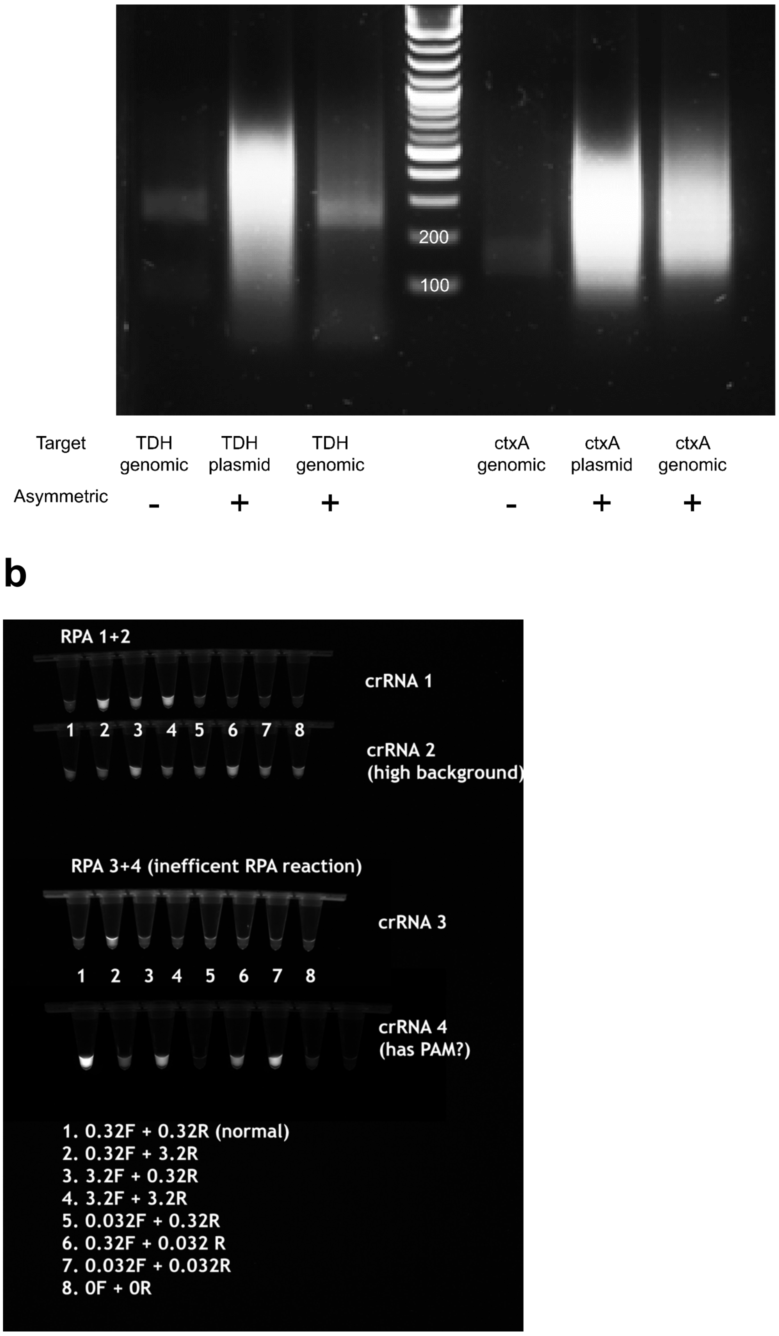


**Supplementary Figure 16. a** Asymmetric RPA of TDH and ctxA plasmid or genomic sample. 0.1ng of plasmid or 10ng of genomic sample was used for the RPA reaction with 0.32μM of primers, and for asymmetric RPA, 10 times molar ratio was utilized either side of primers (3.2μM). The amplified produce was incubated for 20-minutes, and then loaded onto a 2.5% gel stained with SYBR safe for DNA visualization. **b** 2μL of the 20-min asymmetric RPA reaction with 0.1ng plasmid template was added to cas12a reaction mix containing 0.25μM of cas12a-crRNA complex, with 8μM ssDNA-FQ in NEB2.1 (1x) buffer and incubated 20-min at 37-degree Celsius. Note that crRNA1 and crRNA2 is in opposite directions, while crRNA 3 and crRNA 4 is in opposite directions. crRNA is forward facing and hybridizes to the reverse strand of the RPA, same goes for crRNA 3. The opposite is true for crRNA 2 and crRNA 4. crRNA 4 contains a PAM site.

| Components | Cost per kit or tube | Total provided per kit | PECAN usage per reaction (10μL RPA+ 10μL PECAN) | PECAN number of reactions | PECAN Cost (USD) | AIOD CRISPR usage per reaction (25μL AIOD CRISPR) | AIOD CRISPR number of reactions | AIOD CRISPR Cost (USD) |
|---|---|---|---|---|---|---|---|---|
| **ssDNA-FQ** | 325 USD | ~160 nmol | 80 pmol | 2000 | 0.1625 | 200 pmol | 800 | 0.4 |
| **RPA kit** | 160 USD | 5 ml reaction volume | 10 μL | 500 | 0.32 | 25 μL | 200 | 0.8 |
| **LbCas12a** | 264 USD | 2000 pmol | 2 pmol | 1000 | 0.264 | 32 pmol | 62.5 | 4.224 |
| **crRNA** | 350 USD | 50 nmol | 4 pmol | 12500 | 0.028 | 32 pmol | 1562 | 0.22 |
| **T7exo** | 274 USD | 5000 unit | 1 unit | 5000 | 0.055 | / | / | 0 |
| **Total** | | | | | **$0.8** | | | **$5.6** |

**Supplementary Table 5.** Cost analysis of PECAN versus AIOD-CRISPR system with listed usage amount and cost per component.

**Supplementary Table 6.** Jalview visualization of multiple sequence alignment consensus of various gene-targets at the site of RPA primers and crRNA. The search term for the NCBI nucleotide database data mining for the original sequences was also shown.

Search term: vibrio parahaemolyticus tdh (full name)
Sequence length: 0-1000
Manual trim required

| FP alignment | crRNA region | RP alignment |
|---|---|---|
| 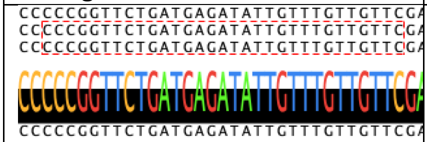 | 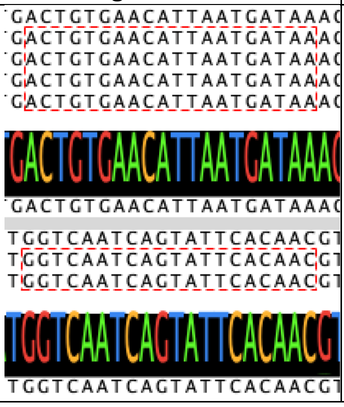 | 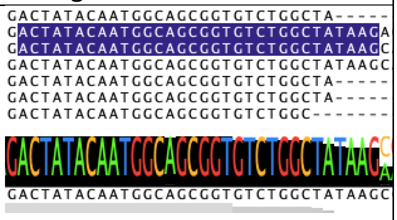 |

Reference genome as template
CP020428.2

Search term: vibrio vulnificus vcgC
Sequence length: 0-1000
No trim required

| FP alignment | crRNA region | RP alignment |
|---|---|---|
| 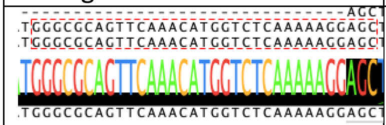 | 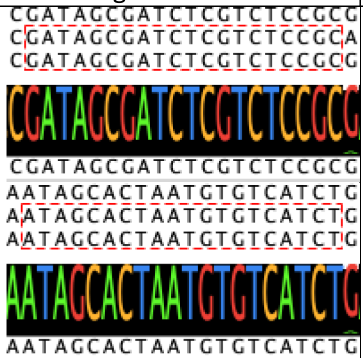 | 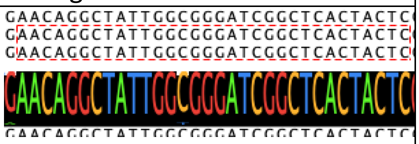 |

Reference genome as template
CP014049.2

Search term: (ctxA) AND "Vibrio cholerae"[porgn:__txid666] NOT (genome shotgun)
Output: 92 sequences
Sequence length: 0 - 1000

| FP alignment | crRNA region | RP alignment |
|---|---|---|
| 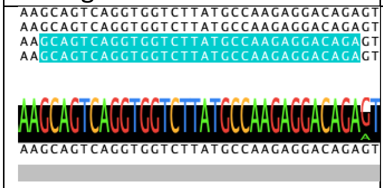 | 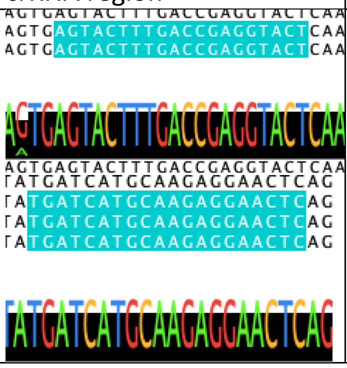 | 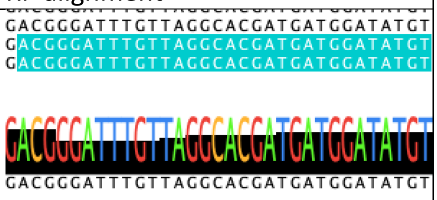 |

Reference genome as template

CP047059

Search term: norovirus GI
Sequence length: 5000-9000
search 2010-2021
Output: 134 sequences

FP alignment

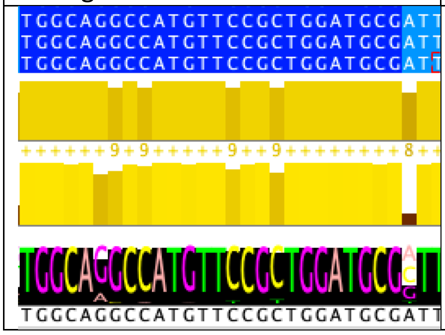


crRNA region

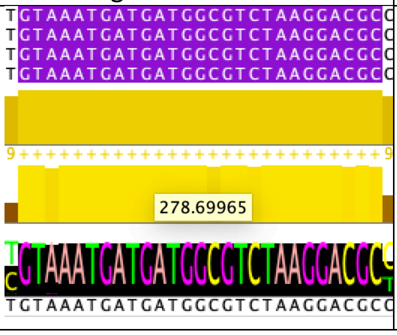


RP alignment

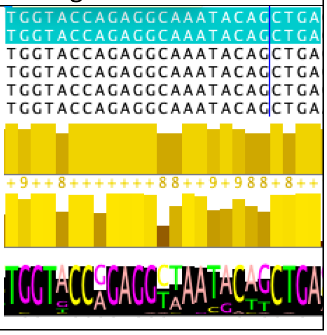


Reference genome as template
MT008453.1 GI :1803118036

**Supplementary Information Reference**

1. Nguyen LT, Smith BM, Jain PK. Enhancement of trans-cleavage activity of Cas12a with engineered crRNA enables amplified nucleic acid detection. Nat Commun. 2020;11(1):4906.
